# Differentiation stage-specific use of cap-independent and cap-dependent translation initiation in hematopoiesis

**DOI:** 10.1101/2025.09.24.678136

**Authors:** Michael C. Mazzola, Ting Zhao, Anna Kiem, Trine A. Kristiansen, Karin Gustafsson, Lai Ping Wong, Emily Scott-Solomon, Marissa D. Fahlberg, Sarah Forward, Emane Rose Assita, Giulia Schiroli, Maris Handley, Youmna Kfoury, Tsuyoshi Fukushima, Samuel Keyes, Azeem Sharda, Jelena Milosevic, Hiroki Kato, Pavel Ivanov, David B. Sykes, Sheldon J. J. Kwok, Ruslan I Sadreyev, Vijay G. Sankaran, Ya-Chieh Hsu, David T. Scadden

## Abstract

Cell stress can increase the use of m^7^G-cap-independent, IRES-mediated translation initiation relative to cap-dependent translation (IRES/Cap). Reporters that quantify IRES/Cap have demonstrated differential activity across cultured cell types and stress conditions. By generating an IRES/Cap reporter mouse, we were able to systematically evaluate IRES/Cap across distinct tissues and cell types during physiological stresses and lineage commitment. Caloric stress invoked the expected boost in IRES/Cap translation regardless of differentiation state, but unexpectedly IRES/Cap progressively increased during hematopoietic and epithelial (hair follicle) differentiation under normal, homeostatic conditions. This was independent of total protein output or cell cycle. Even within cells of a given differentiation state, cells with lower relative-IRES utilization had markedly higher multipotent capability *in vivo*. The RNA processing protein PTBP1 is a mediator of this translation initiation preference. Therefore, low IRES/Cap is a signature of high stemness and suggests modulation of translation initiation participates in cell differentiation state.

## Introduction

Regulation of protein synthesis is a fundamental mechanism by which cells adapt to internal states and external cues (*1*). While transcriptional control shapes long-term gene expression programs, translational control, particularly at the rate-limiting step of initiation, enables rapid and dynamic shifts in protein output (*2*). Under homeostatic conditions, most protein synthesis is driven by cap-dependent translation, which relies on the recognition of a methylated guanosine (m7G) cap at the 5’ end of mRNAs to recruit the eIF4F complex and initiate ribosome loading (*1*). During certain contexts, particularly cellular stress, cap-dependent translation is inhibited, and the contribution of cap-independent protein production is enhanced (*3*). One such mechanism is internal ribosome entry site (IRES)-mediated translation.

IRESes are structured sequences in the UTRs that recruit the pre-initiation complex either directly or through interactions with IRES-transactivating factors (ITAFs), bypassing the need for a 5’ cap (*4*). This mode of translation initiation is best understood in the context of RNA viruses, such as encephalomyocarditis virus (EMCV). Lacking access to the nuclear capping machinery, viral RNAs can harbor IRES elements in their UTRs that co-opt host ribosomes for translation. EMCV also encodes proteases that cleave components of the host’s cap-dependent translation machinery, thereby promoting preferential translation of viral proteins via the IRES-pathway (*5*–*7*).

Beyond viral systems, cap-dependent translation is known to be tightly regulated by signaling pathways, including the nutrient sensor mTOR. mTOR activity influences the availability of the cap-binding protein eIF4E by modulating the phosphorylation of the inhibitory factor 4EBP (*8*). Inhibition of mTOR activity leads to sequestration of eIF4E, suppressing cap-dependent translation while sparing cap-independent translation. Notably, a subset of mammalian mRNAs contains IRES or IRES-like elements that may be selectively translated during stress despite also possessing 5’ caps (*9*).

To assess relative activity of these two modes of initiation, bicistronic reporters are the gold-standard tool, consisting of a single RNA transcript with two open reading frames: the first reporter protein is translated via cap-dependent initiation, and the second reporter protein via an IRES (*10*). Bicistronic reporters enable quantification the IRES/Cap ratio (hereafter referred to as IRES/Cap), providing a readout of cap-independent IRES-mediated translation relative to cap-dependent translation. Because both fluorescent proteins are translated from the same transcript, this approach controls for mRNA abundance and promoter activity, allowing accurate comparisons of IRES/Cap in distinct conditions or cell types. However, studies comparing IRES/Cap between distinct cell types are limited immortalized cell lines (*11, 12*), which inadequately model normal tissue heterogeneity and physiological processes like lineage commitment. While studies have suggested roles for IRES-mediated translation in erythroid progenitor differentiation (*13*) and in embryonic stem cells (*14*), these were restricted to *in vitro* systems and did not systematically assess IRES activity in the context of normal tissue differentiation. Moreover, work in lower eukaryotes has demonstrated that yeast rely on IRES-mediated translation to initiate lineage-specific programs during nutrient starvation, when cap-dependent translation is repressed (*15*), raising the possibility that cap-independent mechanisms might similarly regulate regeneration and cell fate decisions in higher-order eukaryotes.

In this study, we developed the *Translator* mouse, which uses a bicistronic reporter driven by the EMCV IRES to quantify IRES/Cap *in vivo*. In both hematopoietic and epithelial tissues, we found that stem cells exhibit the lowest relative IRES-usage that progressively increases with progenitor lineage commitment. These findings provide a conceptual shift in how IRES-mediated translation is understood: not merely as a stress adaptation, but as a homeostatically regulated process.

### The Translator mouse quantifies IRES-mediated translation relative to canonical cap-dependent translation

To assess the distinct usage of translation initiation in cells at various stages of differentiation, we employed fluorescent bicistronic reporters that permit the simultaneous flow cytometric quantification of IRES/Cap. On a single mRNA transcript, mRFP is translated by cap-dependent translation and GFP by IRES-mediated translation (Fig. 1A). We evaluated the well-characterized EMCV-IRES, as well as the IRES within the 5’ UTR of RUNX1, a transcription factor critical for hematopoiesis and hair follicle differentiation, whose IRES deletion causes embryonic lethality due to defective hematopoietic development (fig. S1, A and E) (*16–18*). HEK293T cells transduced with lentiviral reporters confirmed simultaneous RFP and GFP fluorescence (fig. S1, B and F). To address whether the IRES had cryptic promoter activity, we removed the promoter and the mRFP coding sequence (fig. S1C and G). The absence of GFP fluorescence in promoter-less lentiviral constructs confirmed that neither the EMCV IRES nor the RUNX1 IRES contain cryptic promoters (fig. S1, D and H).

**Fig. 1.**
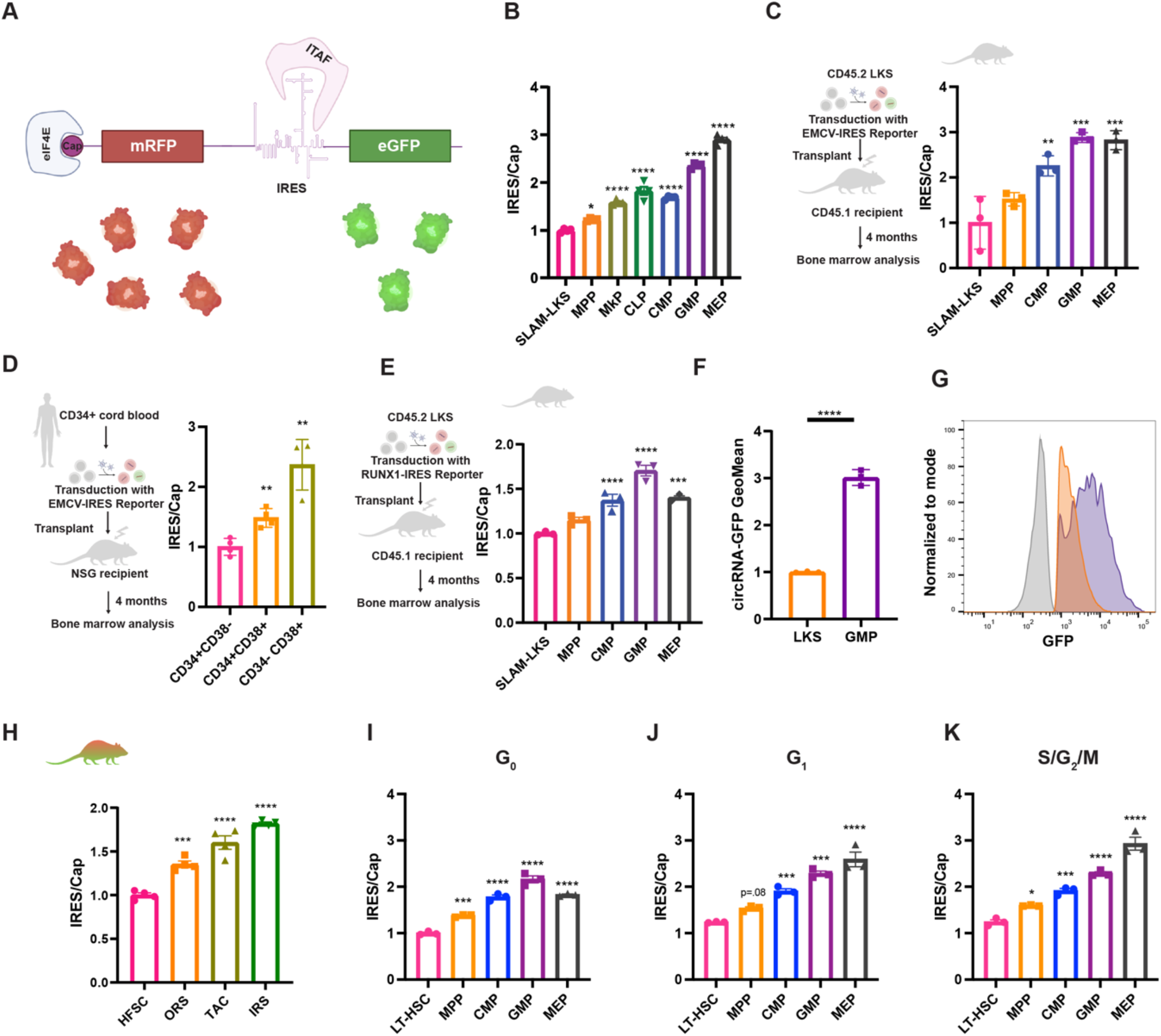
IRES/Cap increases with hematopoietic and hair follicle differentiation. **(A)** Graphical representation of the bicistronic mRNA. mRFP is translated in a cap-dependent manner, and the eGFP is translated by the EMCV-IRES. Assisted by BioRender. **(B)** Analysis of EMCV-IRES/Cap in *Translator* mouse bone marrow HSPCs. Geometric mean of GFP relative to geometric mean of mRFP normalized to the average of SLAM-LKS (*n* = 4 from two independent experiments). Significance is relative to SLAM-LKS. **(C)** Analysis of EMCV-IRES/Cap in 9.5 Gy-lethally irradiated recipients 4-months post-transplantation of LKS transduced with lentivirus encoding bisicstornic reporter. Geometric mean of GFP relative to geometric mean of mRFP normalized to the average of SLAM-LKS (*n* = 3, representative of four independent experiments). Significance is relative to SLAM-LKS. Assisted by BioRender. **(D)** Analysis of EMCV-IRES/Cap in human HSPCs derived from CD34+ cord blood transduced with the EMCV-IRES bicistronic reporter and transplanted in 1.5 Gy-sublethally irradiated NSG recipient mice. Geometric mean of GFP relative to geometric mean of mRFP normalized to the average of CD34+CD38- (n=4, representative of two independent experiments). Significance is relative to CD34+CD38-. Assisted by BioRender. **(E)** Analysis of RUNX1-IRES/Cap in 9.5 Gy-lethally irradiated recipients 4-months post-transplantation of transduced LKS. Geometric mean of GFP relative to geometric mean of mRFP normalized to the average of SLAM-LKS (n=3, representative of two independent experiments). Significance is relative to SLAM-LKS. Assisted by BioRender. **(F)** GFP MFI of cultured LKS and GMP transfected with equal amounts of circRNA encoding GFP and analyzed 24 h post-transfection (n=3, representative of two independent experiments). **(G)** Representative histogram of GFP measured by flow cytometry in LKS and GMP transfected with circRNA encoding GFP. Gray, negative control. **(H)** Analysis of IRES/Cap in *Translator* mouse hair follicles, including hair follicle stem cells (HFSC), outer root sheath (ORS), transit amplifying cells (TACs), and inner root sheet (IRS). Geometric mean of GFP relative to geometric mean of mRFP normalized to the average of HFSC (n=4 from two independent experiments). **(I)** IRES/Cap in *Translator* mouse HSPCs in G_0_, evaluated with single-cell optical barcoding (n=3). **(J)** IRES/Cap in *Translator* mouse HSPCs in G_1_, evaluated with single-cell optical barcoding (n=3). **(K)** IRES/Cap in *Translator* mouse HSPCs in S/G_2_/M, evaluated with single-cell optical barcoding (n=3). Data show individual replicates and mean ± SEM. Significance was assessed using a paired one-way ANOVA **(B-E, H-K)** and a Student’s t test **(F)**. *p ≤ 0.05, **p ≤ 0.01, ***p ≤ 0.001, ****p ≤ 0.0001

To test the fidelity of these reporters, human K562 myelogenous leukemic cell lines expressing these reporters were treated with specific small molecule inhibitors of cap-dependent translation. Rocaglamide inhibits the helicase eIF4A, a critical component of the eIF4F complex, which unwinds secondary structure in the 5’ UTR of mRNAs (*19*). Additionally, torin-2 inhibits mTOR and the availability of the cap-binding protein eIF4E for the cap-binding eIF4F complex formation by reducing dephosphorylation of 4EBP (*8*). As predicted, both compounds significantly increased IRES/Cap by inhibiting cap-dependent translation (fig. S1I).

The *Translator* mouse was generated by pronuclear injection of C57BL/6 zygotes with CRISPR/Cas9 protein and guide RNA to introduce the mRFP-EMCV-IRES bicistronic reporter transgene into the ROSA26 locus (fig. S1J) (*20*). We focused on hematopoiesis since HSPCs have well-defined immunophenotypes and functional assays. *Translator* mouse peripheral blood counts (fig. S1, L-R), HSPC bone marrow (BM) cellularity (fig. S1S), and total BM colony-forming potential (fig. S1T) are comparable to wildtype littermates, confirming that the reporter does not perturb normal hematopoietic differentiation.

### IRES/Cap increases with differentiation

*Translator* mouse BM demonstrated the lowest IRES/Cap in HSCs (defined by SLAM-LKS immunophenotype), with the ratio increasing with HSPC differentiation (Fig. 1B). In similar fashion, IRES/Cap increased with differentiation in HSPCs regenerated from murine HSCs transduced with lentivirus encoding the fluorescent bicistronic reporter and transplanted into lethally irradiated recipients (Fig. 1C). To validate this finding in human cells, we transduced CD34+ cord blood with the reporter and transplanted the transduced cells into immunodeficient mice (Fig. 1D). Consistent with murine HSPC differentiation, IRES/Cap was lowest in human primitive progenitors (CD34+CD38-) compared to differentiated progenitors (CD34-CD38+).

Next, we evaluated a mammalian IRES element from the RUNX1B gene. HSCs were transduced with lentivirus encoding the fluorescent bicistronic reporter of RUNX1B-IRES/Cap translation (Fig. 1E) and transplanted into lethally irradiated recipients. Similar to EMCV-IRES/Cap, the RUNX1B-IRES/Cap was least active in HSCs and highest in GMPs, suggesting that both viral and mammalian IRESes are differentially regulated during early hematopoietic differentiation in mice and humans.

We sought to evaluate IRES-mediated translation rates using a distinct reporter system. Although circRNAs are predominantly known as miRNA sponges and protein decoys (*21*), those being utilized to synthesize proteins must be translated by an IRES since they lack capped 5’-termini (*22*). Transfected circRNA encoding GFP that uses the coxsackievirus B3 (CVB3) IRES translated three-times more GFP in cultured GMPs in comparison to LKS (Fig. 1, F and G). These data demonstrate that this third IRES element is also more active in lineage-committed progenitors.

After demonstrating that IRES/Cap increases with hematopoietic differentiation, we tested whether this finding existed in other tissues. The hair follicle transit amplifying cells (TACs) derived from hair follicle stem cells (HFSCs) rapidly proliferate to produce hair during adult anagen (*23*). Epidermal cells were isolated from *Translator* mice, and the ratio of IRES/Cap was compared between HFSCs, inner root sheath (IRS), TACs, and outer root sheath (ORS), defined by immunophenotype and analyzed by flow cytometry (fig. S2A). As observed in hematopoiesis, IRES/Cap was lowest in the HFSC and increased with hair follicle differentiation (Fig. 1H). IRES/Cap also increased with hair follicle differentiation when IRES/Cap was calculated using immunofluorescent imaging (fig. S2C) of HFSCs (fig. S2D) and TACS (fig. S2E). Taken together, these data confirm that IRES-mediated translation relative to canonical cap-dependent translation is lowest in stem cells and increases with differentiation in at least two regenerative tissues.

Adult HSCs are largely quiescent, while differentiated progenitors actively cycle (fig. S2E) (*24*). Since both cap-dependent and IRES-mediated translation have been shown to be associated with cell-cycle status (*25, 26*), we assessed whether differences in IRES/Cap correlated with differences in cell cycle. Flow cytometric cell cycle analysis requires fixation and permeabilization to stain with DAPI and Ki67 (intracellular), reducing the intensity of mRFP and GFP fluorescence differently across cell types (fig. S2F). To overcome this technical limitation, we employed a single-cell optical barcoding approach using semiconductor laser particles (LPs) (*27, 28*) to measure the same cells with flow cytometry before and after fixation (fig. S2, G and H). IRES/Cap increased with HSPC differentiation, even when controlling for cell cycle status (Fig. 1, I-K), although a larger difference was observed in actively cycling cells.

### IRES/Cap is correlated with clonogenicity and regenerative potential of HSPCs

To assess if IRES/Cap is associated with hematopoietic function, we performed soft agar colony assays, plating HSPCs with the 20% lowest and 20% highest rates of IRES/Cap compared to the average (“mid”) (Fig. 2A). *Translator* mouse SLAM-LKS with low IRES/Cap produced significantly more colonies compared to those with the high IRES/Cap in both primary (Fig. 2B) and secondary (fig. S3A) colony assays. Low IRES/Cap SLAM-LKS colonies were also significantly larger than high IRES/Cap SLAM-LKS (Fig. 2C and fig. S3B). *Translator* mouse GMPs with low IRES/Cap also produced significantly more colonies (Fig. 2D) that were greater in size (Fig. 2E) compared to high IRES/Cap GMPs. Next, we evaluated total rates of protein synthesis in GMPs sorted based on IRES/Cap. GMPs with low IRES/Cap had significantly greater rates of total protein synthesis, as measured by O-propargyl-puromycin incorporation (Fig. 2F). Human cultured HSPCs with low IRES/Cap produced significantly more colonies than those with high IRES/Cap (Fig. 2G). Finally, to evaluate if this trend was specific to the EMCV-IRES, we sorted HSPCs based on RUNX1-IRES/Cap; SLAM-LKS and GMPs with lower utilization of the RUNX1-IRES relative to cap-dependent translation produced more colonies (Fig. 2, H and I). Collectively, low IRES/Cap HSPCs have an increased proliferative and regenerative capacity *ex vivo*.

**Fig. 2.**
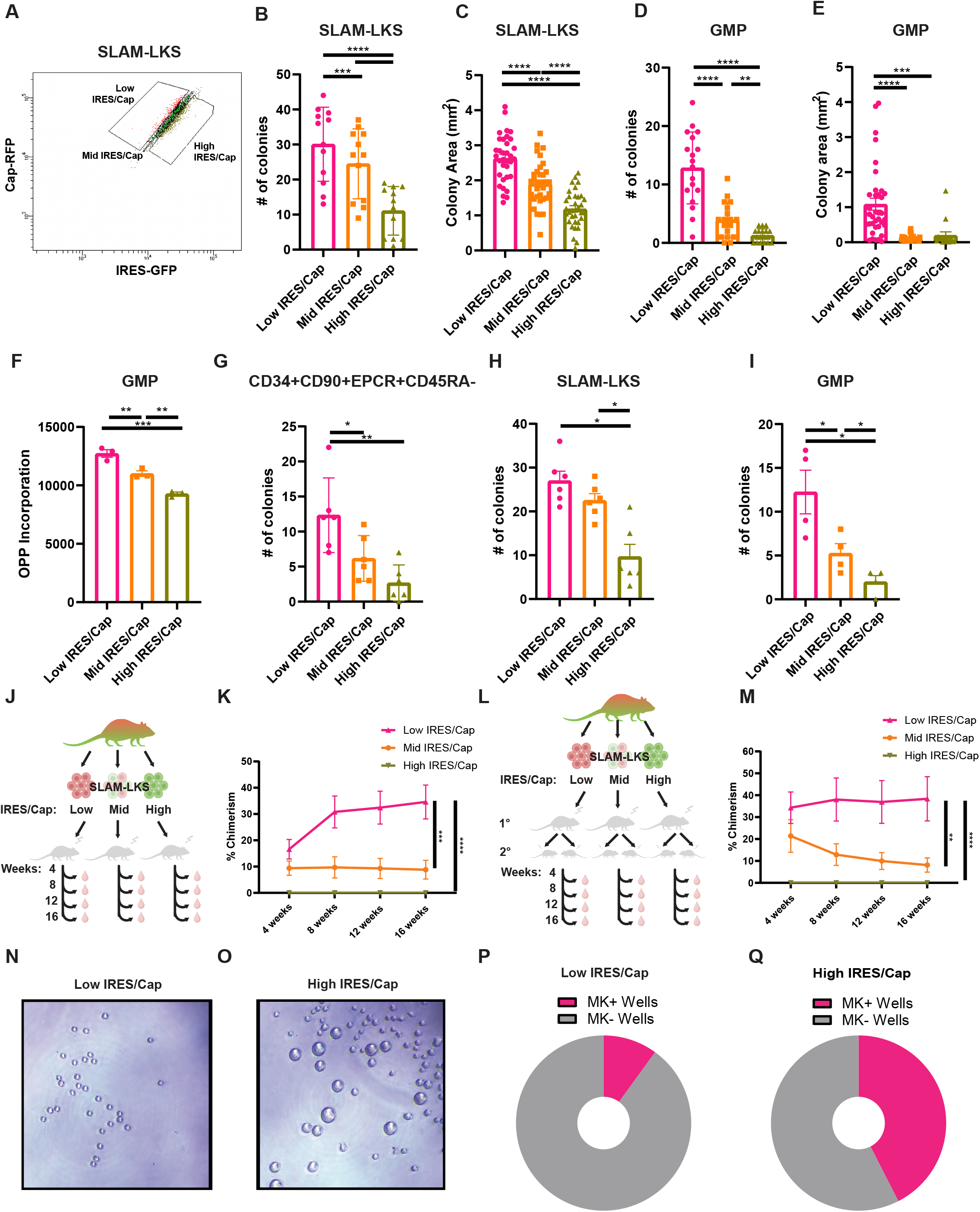
IRES/Cap is associated with clonogenicity and regenerative output of HSPCs. **(A)** Representative gating strategy for sorting low, mid and high IRES/Cap SLAM-LKS by flow cytometry. **(B)** Colony formation of *Translator* SLAM-LKS sorted based on their relative use of IRES/Cap (n=6). **(C)** Colony size (mm^2^) of *Translator* SLAM-LKS sorted based on their relative use of IRES/Cap (n=6). **(D)** Colony formation of *Translator* GMP sorted based on their relative use of IRES/Cap (n=9). **(E)** Colony size (mm^2^) of *Translator* GMP sorted based on their relative use of IRES/Cap (n=9). **(F)** Total protein synthesis rates of *Translator* GMP sorted based on their relative use of IRES/Cap measured by the MFI of AF647 labeled-OPP (n=3). **(G)** Colony formation of CD34+CD90+EPCR+CD45RA-cultured human cord blood transduced with the EMCV-IRES lentiviral bicistronic reporter and sorted based on IRES/Cap (n=3). **(H)** Colony formation of SLAM-LKS based on their relative use of RUNX1-IRES/Cap from transplant recipients (n=3). **(I)** Colony formation of GMPs based on their relative use of RUNX1-IRES/Cap from transplant recipients (n=2). **(J)** Experimental schema for transplantations that compare regenerative potential of *Translator* SLAM-LKS based on IRES/Cap. 20 CD45.2 SLAM-LKS sorted based on IRES/Cap were competed against 200,000 CD45.1 whole bone marrow cells in 9.5 Gy-lethally irradiated. CD45.1 recipients. Peripheral blood was collected retro-orbitally every 4 weeks for 16 weeks. Assisted by BioRender. **(K)** Total chimerism *Translator* SLAM-LKS based on IRES/Cap in primary recipients evaluated every 4 weeks for 16 weeks (n=12). **(L)** Experimental schema for transplantations that compare regenerative potential of *Translator* SLAM-LKS based on IRES/Cap. One million total bone marrow cells were collected from primary recipients (16 weeks post-transplant) and transplanted to 9.5 Gy-lethally irradiated CD45.1 recipients. Peripheral blood was collected retro-orbitally every 4 weeks for 16 weeks. Assisted by BioRender. **(M)** Total chimerism *Translator* SLAM-LKS based on IRES/Cap in secondary recipients evaluated every 4 weeks for 16 weeks (n=12). **(N)** Representative image of culture of a single low IRES/Cap SLAM-LKS after 5 days of culture. **(O)** Representative image of culture of a single high IRES/Cap SLAM-LKS after 5 days of culture. **(P)** Pie chart depicting the percentage of wells with megakaryocytes (Mk) after 5 days of culturing a single low IRES/Cap *Translator* SLAM-LKS (n=20). **(Q)** Pie chart depicting the percentage of wells with Mks after 5 days of culturing a single high IRES/Cap *Translator* SLAM-LKS (n=20). Data show individual replicates and mean ± SEM. *p ≤ 0.05, **p ≤ 0.01, ***p ≤ 0.001, ****p ≤ 0.0001. Significance was assessed using a two-way ANOVA with Dunnett’s One-way ANOVA **(B-I)** or an ANOVA with Tukey’s multiple comparisons test **(K & M)**.

To evaluate the impact of IRES/Cap on HSC function *in vivo*, we transplanted SLAM-LKS based on IRES/Cap in competition with CD45.1 marrow into lethally irradiated recipients (Fig. 2J). Low IRES/Cap SLAM-LKS resulted in significantly greater peripheral blood chimerism over 16-weeks across all lineages compared to average (mid) IRES/Cap and high IRES/Cap (Fig. 2K and fig. S3, C-E). The self-renewal of LT-HSC defined by SLAM-LKS based on IRES/Cap was evaluated by performing serial transplantation (Fig. 2L). Low IRES/Cap SLAM-LKS had persistent chimerism in secondary recipients, whereas mid IRES/Cap had reduced regenerative potential after 16-weeks (Fig. 2M and fig. S3, F-H).

To investigate a potential reason behind the low chimerism observed from high IRES/Cap SLAM-LKS, we sorted SLAM-LKS based on IRES/Cap (fig. S3I) and grew them in HSC expansion liquid cultures. After five days of culture, high IRES/Cap SLAM-LKS produced significantly more megakaryocytes than low IRES/Cap (Fig. 2, N-Q).

Collectively, these data indicate that IRES/Cap distinguishes functional subsets of HSC (SLAM-LKS). Low IRES/Cap SLAM-LKS have the greatest capacity for long-term reconstitution of blood in serial transplantations and high IRES/Cap SLAM-LKS are megakaryocyte-primed. Therefore, the relative use of these translation initiation mechanisms is indicative of functional distinctions between immunophenotypically identical cells.

### IRES utilization is associated with the expression of differentiation-related genes

Since IRES/Cap correlates with differentiation-status within a particular immunophenotype, we evaluated the transcriptome of HSPCs sorted based on their relative use of IRES-mediated translation. Translator SLAM-LKS, GMP and MEP gated based on the ratio of IRES/Cap were evaluated by single-cell transcriptomic analysis (100 cells/condition, Fig. 3, A-F, Table S1). Genes associated with megakaryocyte lineage priming were upregulated in high IRES/Cap SLAM-LKS (*29, 30*), compared to low IRES/Cap SLAM-LKS, which had greater expression of genes associated with stemness, including Fdg5, Mllt3, and Procr (Fig. 3G) (*31–33*), CD41 protein, a cell-surface marker of megakaryocyte-priming (*34*), was quantified in SLAM-LKS. High IRES/Cap SLAM-LKS had significantly greater CD41 protein compared to those with low IRES/Cap (Fig. 3, H and I), which is consistent with the lower chimerism of high IRES/Cap SLAM-LKS in competitive transplant experiments (Fig. 2K) and high megakaryocytic output *ex vivo* (Fig. 2, N-Q). Finally, low IRES/Cap SLAM-LKS had significantly greater expression of genes included in Hallmark 2020 terms associated with inflammation, specifically interferon-γ signaling, which is known to be expressed in HSCs (fig. S4, A and B) (*35*).

**Fig. 3.**
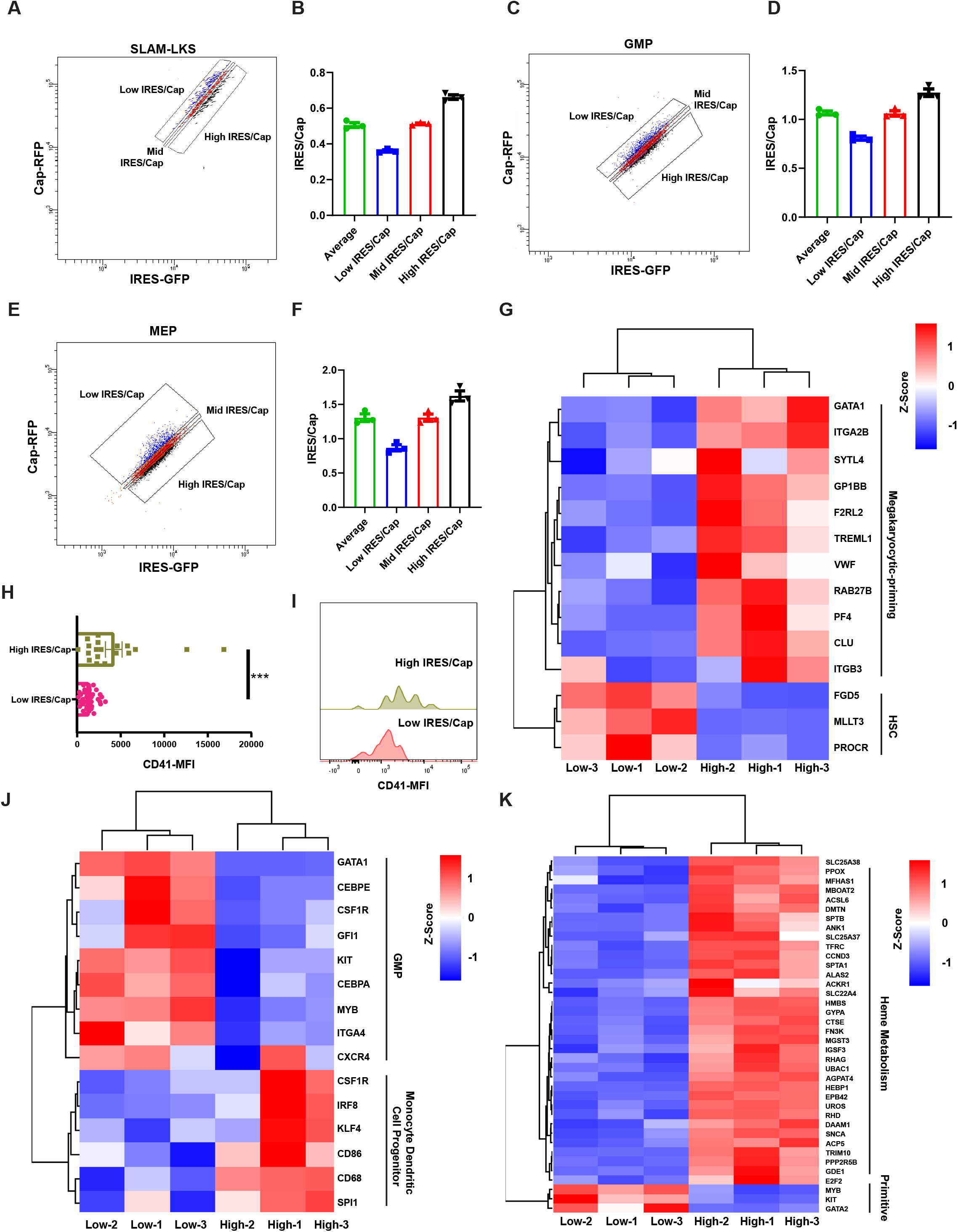
IRES/Cap is associated with differentiation gene expression. **(A)** Representative gating strategy for sorting SLAM-LKS based on IRES/Cap by flow cytometry. **(B)** Quantification of IRES/Cap based on gating strategy in **(A)**. IRES/Cap is normalized to that of the entire population of SLAM-LKS (n=3). Data show mean ± SEM. **(C)** Representative gating strategy for sorting GMP based on IRES/Cap by flow cytometry. **(D)** Quantification of IRES/Cap based on gating strategy in **(D)**. IRES/Cap is normalized to that of the entire population of GMP (n=3). Data show mean ± SEM. **(E)** Representative gating strategy for sorting MEP based on IRES/Cap by flow cytometry. **(F)** Quantification of IRES/Cap based on gating strategy in **(G)**. IRES/Cap is normalized to that of the entire population of MEP (n=3). Data show mean ± SEM. **(G)** DEGs enriched in high IRES/Cap SLAM-LKS compared to low IRES/Cap SLAM-LKS (n=3). **(H)** CD41-MFI on index-sorted low and high IRES/Cap *Translator* SLAM-LKS measured by flow cytometry. **(I)** Representative histogram of CD41-BV711 on index-sorted low and high IRES/Cap *Translator* SLAM-LKS measured by flow cytometry. **(J)** DEGs enriched in high IRES/Cap GMP compared to low IRES/Cap GMP (n=3). **(K)** DEGs enriched in high IRES/Cap MEP compared to low IRES/Cap MEP (n=3). Data show individual replicates and mean ± SEM. Significance was assessed using a Student’s t test **(H)**. ***p ≤ 0.001

Differentially expressed genes enriched in high IRES/Cap GMPs were functionally associated with monocyte dendritic cell progenitors (Csf1R, Irf8, Flg4, CD86, CD68, Spi1) (Fig. 3J and fig. S4D), consistent with myeloid differentiation (*36–38*). Conversely, low IRES/Cap GMPs had higher expression of genes critical for the myeloid progenitor state (Gata1, Cebpe, Csf1r, Gfi1, Kit, Cebpa, Myb, Itga4, Cxcr4) (Fig. 3J and fig. S4C) . Similarly, high IRES/Cap MEPs had significantly greater expression of genes involved in heme metabolism compared to low IRES/Cap, suggesting IRES/Cap is positively correlated with MEP differentiation (Fig. 3K and fig. S4, E and F). These transcriptomic analyses collectively support our findings that HSPCs with low relative IRES utilization are more primitive and poised for high cell output, whereas those with high relative IRES utilization are more differentiated and perform effector functions.

### Relative IRES usage is positively regulated by PTBP1

ITAFs are capable of regulating IRES-mediated protein production (*4*). To identify specific regulators of IRES/Cap in hematopoietic cells, we performed a genome-wide CRISPR interference (CRISPRi, dCas9-KRAB) screen on K562 cells expressing the EMCV-IRES bicistronic reporter (Fig. 4A, Table S2) . The dCas9 activity was validated using guides to known targets (fig. S4G). Following transduction with a pooled guide RNA library, K562 cells were sorted on day 5 based on the ratio of IRES/Cap, and gRNA were sequenced. Putative positive regulators of IRES/Cap (i.e., KEGG terms associated with gRNA enriched in low IRES/Cap) were ribosome biogenesis regulators (Fig. 4B), whereas putative negative regulators were enriched for RNA transport proteins (Fig. 4C). These findings implicated ribosome abundance as a key regulator of differential IRES translation in HSPCs.

**Fig. 4.**
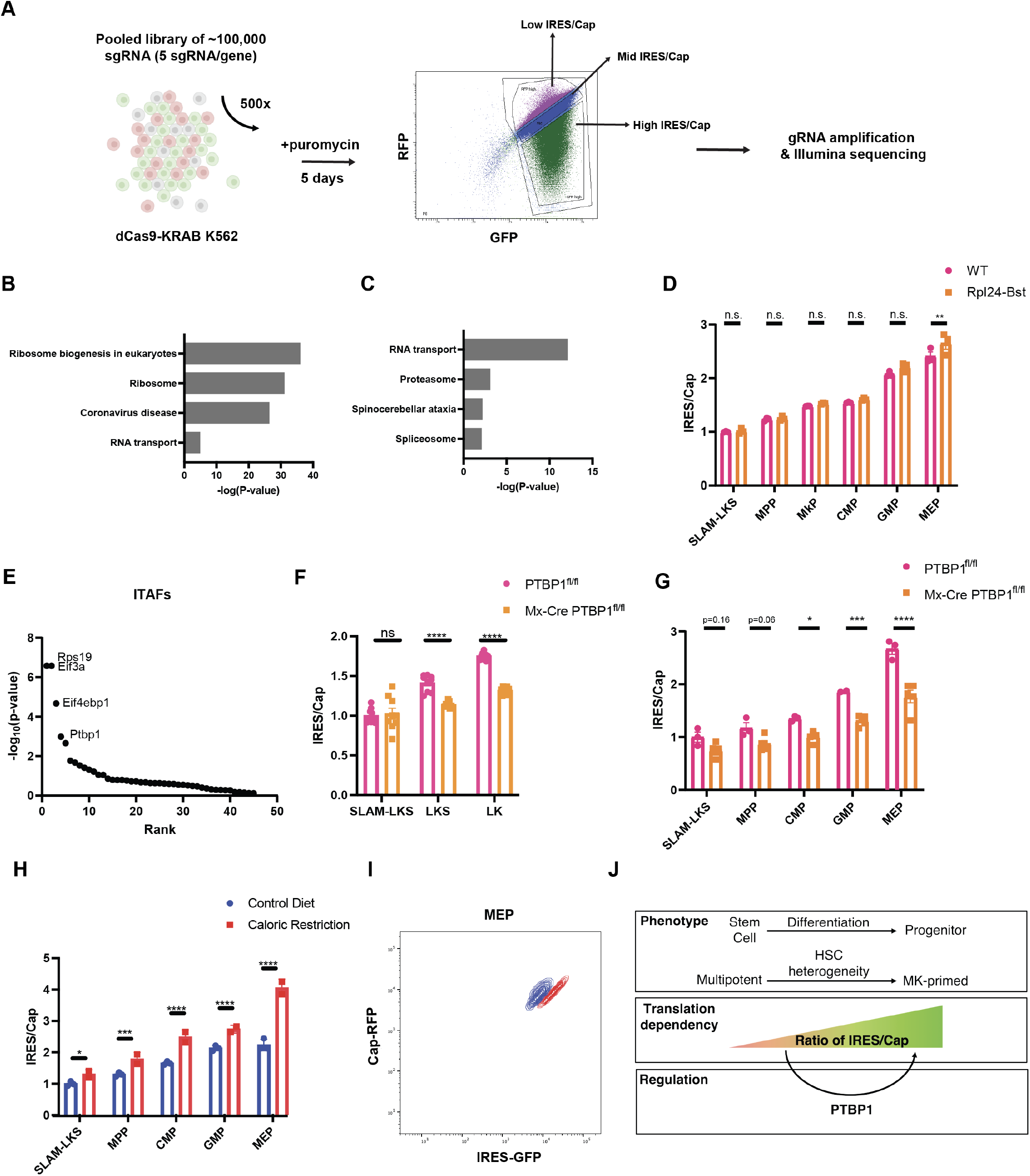
IRES/Cap is regulated by PTBP1 and caloric restriction. **(A)** Graphical representation of a genome-wide CRISPR interference screen workflow to identify positive and negative regulators of IRES/Cap. Assisted by BioRender. **(B)** Top KEGG-terms associated with genes targeted by sgRNAs enriched in “low IRES/Cap” gate (i.e. putative positive regulators of IRES/Cap) (n=3). **(C)** Top KEGG-terms associated with genes targeted by sgRNAs enriched in “high IRES/Cap” gate (i.e. negative regulators of IRES/Cap) (n=3). **(D)** Analysis of EMCV-IRES/Cap in *Translator* Rpl24-Bst HSPCs compared to WT littermate controls measured by flow cytometry (n=3). **(E)** Known IRES-transactivating factors (*4*) ranked in order of significance as putative positive regulators of IRES/Cap in K562 (n=3). **(F)** EMCV-IRES/Cap in PVA-expanded PTBP1^fl/fl^ and Mx-Cre PTBP1^fl/fl^ HSPCs measured by flow cytometry (n=4). **(G)** EMCV-IRES/Cap in PTBP1^fl/fl^ and Mx-Cre PTBP1^fl/fl^ HSPCs regenerated from transduced LKS in CD45.1 recipients. (n=3 for PTBP1^fl/fl^ and n=5 for Mx-Cre PTBP1^fl/fl^). **(H)** Analysis of EMCV-IRES/Cap in *Translator* mouse HSPCs for animals fed with control diet (n=3) or those that underwent caloric restriction (n=2). Geometric mean of GFP relative to geometric mean of mRFP normalized to the average of SLAM-LKS. Significance is relative to SLAM-LKS from control diet. **(I)** Representative flow plot of MEPs analyzed from mice that were fed normal diet or underwent caloric restriction. **(J)** Illustration depicting the key findings that IRES utilization relative to cap-dependent translation (IRES/Cap) increases with differentiation, which is partially regulated by PTBP1. Data show individual replicates and mean ± SEM. Significance was assessed using a two-way ANOVA **(D, F-H)**. *p ≤ 0.05, **p ≤ 0.01, ***p ≤ 0.001, ****p ≤ 0.0001

Stem cells produce less total protein than more differentiated progenitors (*39–42*). Based on the CRISPRi screen results, we questioned if IRES/Cap reflects differences in total rates of protein synthesis, rather than differentiation status. To address this question, we crossed the translator mouse to the Rpl24^Bst/+^ model, which has well-characterized defects in ribosome biogenesis, slower rates of total protein synthesis in HSPCs, and impaired HSC function (*39*). IRES/Cap was comparable between WT and Rpl24^Bst/+^ HSPCs, suggesting that perturbation of total translation rates by interfering with ribosome function does not alter IRES/Cap (Fig. 4D).

We evaluated how known ITAFs influenced the ratio of IRES/Cap (Fig. 4E) (*4*). Ranking fourth, PTBP1 inhibition reduced IRES/Cap. PTBP1 is a known regulator of ribosomal biogenesis and a stabilizer of IRES elements containing the CUUU motif (*43*), which is present in both the EMCV and RUNX1 IRESes. Blood specific deletion of PTBP1^fl/fl^ by Mx-Cre impairs red blood cell (RBC) production and HSC function in serial transplantations (*44*), suggesting that PTBP1 functionally regulates hematopoietic differentiation.

We cultured and transplanted Mx-Cre PTBP1^fl/fl^ and PTBP1^fl/fl^ HSCs transduced with the EMCV-IRES reporter and evaluated the ratio of IRES/Cap after five days of liquid culture and four months post-transplantation. IRES/Cap is lower in more differentiated Mx-Cre PTBP1^fl/fl^ HSPCs *in vitro* (Fig. 4F) and *in vivo* (Fig. 4G) compared to wildtype PTBP1^fl/fl^. This finding was replicated from Translator-iCAS9 HSPCs transduced with PTBP1- or non-targeting guides and transplanted into lethally irradiated recipients (fig S4, H and I). Therefore, PTBP1 is one of multiple critical component of the IRES/Cap regulatory mechanisms in hematopoiesis. It is highly likely, however, that additional ITAFs must be responsible for the increase in IRES/Cap with differentiation, since PTBP1^-/-^ differentiated progenitors do not have equivalent IRES/Cap to WT HSCs.

### Caloric restriction increases IRES/Cap in both stem and progenitor cells

IRES-translation persists when mTOR signaling is pharmacologically inhibited or reduced in conditions of nutrient deprivation (*45*). In line with this observation, IRES/Cap increased in each subset of hematopoietic cells in the setting of calorie restriction (Fig. 4, H and I) or short-term fasting (fig. S4J). These data parallel the effect of chemical inhibitors of cap-dependent translation in K562 cells (fig. S1I), and suggest that while IRES utilization is developmentally constrained in hematopoiesis, it can be activated under stress. Calorie restricted animals also had significant reductions in RBC and white blood cell (WBC) counts, though the platelet count was preserved (fig. S4, K-M). Ultimately, we propose that IRES utilization is enhanced not only during stress, but also as a critical signature of normal differentiation and homeostasis.

## Discussion

The *Translator* mouse revealed that HSCs had the lowest IRES/Cap ratios, and that this ratio increased with differentiation. Manipulation of total translation rates, such as through the Rpl24^Bst/+^ genotype, was insufficient to alter IRES/Cap ratios across hematopoiesis. Collectively, these data indicate that cap-dependent translation machinery and ITAFs, like PTBP1, are regulated distinctly in differentiating tissues, influencing relative IRES utilization independently of overall translation rates (Fig. 4J).

Since IRES-mediated translation is differentially utilized throughout hematopoietic and hair follicle differentiation, it is likely that this mechanism plays a broader role in other stem cell-dependent contexts. Furthermore, the relative use of IRES correlates with functional heterogeneity within immunophenotypically identical cells. IRES/Cap measured by the *Translator* mouse bicistronic reporter identifies a subset of SLAM-LKS capable of long-term regenerative output. Strikingly, higher IRES/Cap among immunophenotypic HSCs associates with accelerated differentiation and a complete failure to function as stem cells. Therefore, IRES use negatively correlates with stemness and may serve as a capacitor for differentiation.

In murine hematopoietic differentiation, translation regulation is critical to the activation of stem-like megakaryocytic progenitors in response to acute inflammation (*30*), and RUNX1 has been shown to be a critical regulator of megakaryocyte-primed HSC differentiation (*46*). Since RUNX1B protein expression is regulated in part by an IRES, these data may explain why high IRES/Cap SLAM-LKS produced significantly more megakaryocytes *ex vivo* and very low chimerism in primary and secondary transplant recipients. Additionally, HSPCs with greater expression of RUNX1B protein have lower clonogenicity (*47*), as we observed with HSPCs with greater RUNX1B-IRES activity. Regulators of erythroid-differentiation Bag1 and Csde1 contain putative IRES-elements in their 5’UTRs that are regulated by Rps19, an ITAF that scored significantly as a positive regulator of IRES/Cap in K562 (*13*). Collectively, these data suggest that more differentiated progenitors, particularly along the megakaryocytic and erythroid lineages, utilize IRES-mediated protein production to enable differentiation and cell production. However, quantification of endogenous IRES-mediated protein production is technically infeasible with methods such as ribosome profiling, given that all mRNAs are capped (including those with IRESes) and the coding-sequence of IRES-containing isoforms is highly homologous with non-IRES containing isoforms.

PTBP1 has been associated with enhancing virus production, including integrating viruses like human immunodeficiency virus (HIV) (*48*) and non-integrating viruses like Dengue virus (*49*), which both have reported IRES elements (*50–53*). Therefore, low stem cell expression of PTBP1 may protect them from producing viral proteins that depend on IRES-translation. Given their self-renewing capabilities, stem cells may constrain the translation of IRES-mediated viral proteins to protect themselves from durable genomic modifications that pose oncogenic risk. HSC infection by non-integrating RNA viruses, like Dengue Virus, has been shown to impair the regenerative capacity of transplanted HSCs lacking intact interferon-signaling (*35*). Since stem cells have low tropism to many viruses as a mechanism of protection, it is challenging to evaluate if lower viral IRES-mediated translation is an additional layer of protection. However, it is notable that human HSCs have been shown to be resistant to HIV infection despite documented receptor and co-receptor expression (*54, 55*).

Furthermore, expression of endogenous retroviral genetic elements has recently been demonstrated to be important in HSC response to stress (*56*). The expression of those likely triggers an interferon response that may be adaptive, but translation of retroviral proteins to enable further integration could be problematic for HSC genomic stability. It is intriguing to consider whether the restriction on IRES mediated translation in stem cells serves as a means of preserving stem cell genetic integrity.

## Materials and Methods

### Mice

All animal experiments were approved by the Institutional Animal Care and Use committee at Massachusetts General Hospital (IACUC protocol #2016N000085). Mice were housed in a temperature- and humidity-controlled environment with a 12-h light-12-h dark cycle and food and water ad libitum. Wild-type CD45.2 C57BL/6J (strain #000664), CD45.1 (B6.SJL-Ptprc < a > Pepc < b> /BoyJ, strain #002014), KH2/iCas9 (B6;129S4-*Gt(ROSA)26Sor*^*tm1(rtTA*M2)Jae*^ *Col1a1*^*tm1(tetO-cas9)Sho*^/J, strain #029415), Rpl24^Bst^ (C57BLKS-Rpl24^Bst^/J, strain #000516), and NSG mice (NOD.Cg-PrkdcscidIl2rgtm1Wjl/SzJ, strain #005557) were purchased from the Jackson Laboratory. Mice were gender matched and 8-14 weeks of age in all experiments unless stated otherwise.

### Generation of the Translator mouse

The Translator mouse was generated as previously described (*20*) by the Harvard Genome Modification Facility. The knock-in construct was modified from pR26CAG/GFP Dest (#74286, Addgene) by VectorBuilder to include the bicistronic fluorescent reporter (mRFP-EMCV IRES-eGFP) downstream of a CAG promoter followed by a puromycin-transcription stop cassette flanked by Loxp sites. The puromycin-transcription stop cassette was removed by an in vitro Cre Recombinase reaction (#M0298, New England BioLabs), per the manufacturer’s instructions. The donor DNA consists of a 1,083bp left homology arm and a 4,341bp right homology arm to target to the ROSA26 locus. Pro-nuclear injection of the donor DNA paired with a purified Cas9 Nuclease V3 protein (#1081058, Integrated DNA Technologies) and a gRNA targeting ROSA26 (ACUCCAGUCUUUCUAGAAGA) was performed on C57BL/6J zygotes. The transgenic progenies were evaluated for cassette integration by flow cytometry on peripheral blood by retro-orbital bleeding using heparinized micro-hematocrit capillary tubes (#22-362-566, Fisherbrand) and depleted of red blood cells by ACK lysis (#18-156-721, Quality Biological). Heterozygous transgene positive animals were bred with wild-type CD45.2 C57BL/6J mice purchased from the Jackson Laboratory.

### Flow cytometry analysis and sorting

Mice were sacrificed by CO2 asphyxia. For hematopoietic studies, total bone marrow was collected by crushing the tibias, femurs, hips, humeri, and spine. Lineage positive cells were depleted with a lineage cell depletion kit (#130-090-858, Miltenyi Biotec), as per the manufacturer’s instructions. Lineage depleted marrow was stained in PBS supplemented with 2% FBS using the following antibodies: Sca1-BV785 (Clone D7, #108139 Biolegend, 1:200), cKit-APC-eFluor 780 (Clone 2B8, #47-1171-82, Invitrogen, 1:100), CD150-PE-Cy7 (Clone TC15-12F12.2, #115914, Biolegend, 1:200), CD48-BUV737 (Clone HM48-1, #749666, BD, 1:200), CD16/32-BV605 (Clone 2.4G2, #563006, BD, 1:200), CD34-AF647 (Clone RAM34, #560230, BD, 1:30), CD41-BV711 (Clone MWReg30, #740712, BD, 1:200), CD127-eFluor 450 (Clone A7R34, #48-1271-82, eBiosciences, 1:100), biotinylated lineage cocktail comprising of equal parts (CD8a (Clone 53-6.7, #553029, BD), CD3ε (Clone 145-2C11, #553060, BD), CD45R (Clone RA3-6B2, #553086, BD), GR1 (Clone RB6-8C5, #553125, BD), CD11b (Clone M1/70, #553309, BD), Ter119 (Clone Ter-119, #553672, BD), CD4 (Clone GK1.5, #553728, BD) (1:50) followed by Streptavidin AF700 conjugate (#S21383, Invitrogen, 1:300). Dead cells were excluded from analysis by staining with DAPI (#62248, Thermo Scientific, 0.1 µg/mL). The following cell-surface markers were used to define: SLAM-LKS (DAPI Lin-Sca1+cKit+CD150+CD48-), MPP (DAPI Lin Sca1+cKit+CD150-CD48+), LKS (DAPI Lin Sca1+cKit+), MkP (DAPI Lin Sca1-cKit+CD150+CD41+), CMP (DAPI Lin Sca1-cKit+CD34+CD16/32-), CLP (DAPI Lin Sca1medcKitmedCD127+), GMP (DAPI Lin Sca1-cKit+CD34+CD16/32+), and MEP (DAPI Lin Sca1-cKit+CD34-CD16/32-). Marrow was stained for 1.5 hours for primary antibodies and 15 minutes for Streptavidin secondary stains at 4 °C. For analysis and sorting of human cord blood transplantations into NSG mice, marrow was depleted of RBCs by performing ACK lysis (#18-156-721, Quality Biological) and stained in PBS supplemented with 2% FBS using the following antibodies: hCD19-BV711 (Clone HIB19, #302246, BioLegend, 1:100), hCD3-BV711 (Clone UCHT1, #300463, BioLegend, 1:100), hCD34-APC-Cy7 (Clone 581, #343514, BioLegend, 1:100), hCD38 PE-Cy7 (Clone HIT2, #303516, BioLegend, 1:100), hCD90-BV421 (Clone 5E10, #328122, BioLegend, 1:50), and hCD45RA-BUV737 (Clone HI100, #564442, BD, 1:100). For analysis and sorting of human cord blood for colony assays, cultured and transduced CD34+ cells were stained with hCD34-APC-Cy7 (Clone 581, #343514, BioLegend, 1:100), hCD45RA-BUV737 (Clone HI100, #564442, BD, 1:100), hCD90-BV711 (Clone 5E10, #328139, BioLegend, 1:50), and hCD201-APC (Clone RCR-401, #351906, BioLegend, 1:25). Dead cells were excluded from analysis by staining with DAPI (#62248, Thermo Scientific, 0.1 µg/mL).

For hair follicle analysis, translator mice in second anagen (P29-P30) were scarified by CO2 asphyxia. Mouse back skin was dissected, and the fat layer was scraped using a surgical scalpel (#29550, Exel Int). The skin was incubated on an orbital shaker (75 rpm) dermal-side down in Collagenase (0.25%, Sigma) diluted in Hanks’ Balanced Salt Solution for 1 h at 37 °C. Dermal cells were obtained by scraping the dermal side with a dulled surgical scalpel. Dermal cells were then spun at 500g at 37 °C for 5 minutes and resuspended in 0.25% Trypsin-EDTA (#25200072, ThermoFisher Scientific) and incubated at 37 °C for 30 minutes with agitation. Dermal single-cell suspensions were obtained by filtering trypsin suspension through 70-µm and 40-µm filters. For epidermal cells, the post-Collagenase digested skin was incubated on an orbital shaker (75 rpm) dermal-side down in 0.25% Trypsin-EDTA (#25200072, ThermoFisher Scientific) and incubated at 37 °C for 30 minutes. Epidermal single-cell suspensions were obtained by scraping the epidermal side with a dulled surgical scalpel and filtering through 70-µm and 40-µm filters. Dermal and epidermal cells were combined, blocked with CD16/32 Fc (#553142, BD) and incubated for 15 minutes at 4 °C. Cells were incubated with anti-CD140a-Biotin (eBiosciences 13-1401-82, 1:200) for 30 minutes at 4 °C. CD140a+ cells were depleted using Dynabeads M-280 Streptavidin (#11205D, Invitrogen). Cells were incubated with CD49f-APC-Integrin α6 (#313616 Biolegend, 1:500), CD45.2-APC-Cy7 (#109824 Biolegend, 1:250), Sca1-BV785 (#108139 Biolegend, 1:200), and CD34-BV421 (#562608 BD, 1:50). Dead cells were excluded from analysis by staining with DAPI (#62248, Thermo Scientific, 0.1 µg/mL). The following cell-surface markers were used to define: HFSC (DAPI CD45.2-CD49f+CD34+Sca1-), outer root sheath (ORS, DAPI CD45.2-CD49fhiSca1-), transit amplifying cells (TACs, DAPI CD45.2-CD49f^mid^CD34-Sca1-), and inner root sheath (IRS, DAPI CD45.2-CD49f^lo^CD34-Sca1-). All cytometry analysis above was performed using FlowJoTM software version 10.9.0.

When evaluating the ratio of IRES/Cap, the geometric mean fluorescence intensity (geo MFI) was calculated on reporter positive cells for FITC (GFP) and PE-Texas Red (mRFP1) using FlowJo and then divided. When sorting HSPCs based on IRES/Cap, the ratio of was calculated for the entire population of interest. Then gates were drawn to include approximately 20% of cells with the lowest IRES/Cap, 20% of with the median IRES/Cap (equal to IRES/Cap of the parent population), and 20% with the highest IRES/Cap.

### Cell Cycle Analysis

Bone marrow was lineage depleted and HSPCs were immunophenotyped as described above. Lineage depleted marrow was stained with Sca1-PE-Cy5 (Clone D7, #108109 Biolegend, 1:200), cKit-BV650 (Clone ACK2, #752697, BD Biosciences, 1:100), CD150-PE-Cy7 (Clone TC15-12F12.2, #115914, Biolegend, 1:200), CD48-PerCP-Cy5.5 (Clone HM48-1, #103421, BioLegend, 1:200), CD16/32-BV605 (Clone 2.4G2, #563006, BD Biosciences, 1:200), CD34-AF647 (Clone RAM34, #560230, BD, 1:30), and the aforementioned lineage cocktail, followed by Streptavidin-AF700 conjugate (#S21383, Invitrogen, 1:300). Prior to fixation and permeabilization, live marrow samples were barcoded with biotin-coated LPs as described previously (*1*) via biotinylated CD45 (Clone 30-F11, #103104, BioLegend), CD105 (Clone MF7/18, #13-1051-85, eBioscience), CD150 (Clone TC15-12F12.2, #115908, BioLegend), CD41 (Clone MWReg30, #133930, BioLegend), and H-2Kb/H2-Db (Clone 5041.16.1, #PIMA517998, Invitrogen) and purified streptavidin (#405151, BioLegend).. Afterwards, samples were stained for viability, acquired & captured on a multi-pass flow cytometer (LASE Innovation Inc.). The barcoded bone marrow samples were then fixed and permeabilized using the BD Cytofix/CytopermTM Fixation/Permeabilization Kit as described above and stained for AF555-Ki67 (Clone B56, #558617, BD Biosciences, 1:10) and DAPI. In the second cycle’s measurement, 3 biological replicates were acquired at 10 µL/min to maximize DAPI resolution.

Single-color compensation controls were acquired the same day with UltraComp eBeadsTM, ArCTM beads, and GFP BrightComp eBeadsTM (Invitrogen) at gain settings dictated by instrument voltration. ∼7,500 events per compensation control sample were acquired to generate the compensation matrix used for each cycle. For each sample analyzed with the multi-pass method, the FCS files from both cyclic measurements (pre- and post-fix/perm) were concatenated by LP barcode by the LP matching algorithm as described previously (Kwok et al., 2022). The full compensation matrix was manually constructed by concatenating 2 copies of the original 12 x 12 compensation matrix into a 24 x 24 element matrix where elements corresponding to fluorophores of markers measured in different cycles were assigned to 0. When sorting HSPCs based on IRES/Cap, the ratio was calculated for the entire population of interest. The positive Ki67 gate was distinguished by the Alexa Fluor® 555 isotype control (#9641, Cell Signaling Technologies, 1:20), which was stained using the same antibodies as the samples, except using anti-IgG1 Alexa Fluor® 555 in place of Ki67 Alexa Fluor® 555.

### Culture of murine LKS and GMP

GMPs were sorted as described from WT (C57Bl6/J) and cultured in StemSpan™SFEMII (Stem Cell Technologies) supplemented with penicillin (#15140-122, Gibco, 100 IU/mL), streptomycin (#15140-122, Gibco, 100 IU/mL), glutamine (#25030081, Gibco, 2 mM) in addition to mouse recombinant cytokines: SCF (#250-03, Peprotech, 100 ng/mL), IL3 (#213-13, Peprotech, 20 ng/mL), and IL6 (#216-16, PeproTech, 20 ng/mL). LKS were sorted as described above and cultured in StemSpan™SFEMII (Stem Cell Technologies) supplemented with penicillin (#15140-122, Gibco, 100 IU/mL), streptomycin (#15140-122, Gibco, 100 IU/mL), glutamine (#25030081, Gibco, 2 mM) in addition to mouse recombinant cytokines: SCF (#250-03, Peprotech, 100 ng/mL), IL3 (#213-13, Peprotech, 20 ng/mL), TPO (#315-14, PeproTech, 50 ng/mL), and FLT3-ligand (#250-31L, PeproTech, 100 ng/mL).

### In Vitro Protein Translation Assay

GMPs sorted based on the ratio of IRES/Cap from lineage depleted Translator mouse bone marrow were incubated in a humidified 37°C incubator for 30 minutes in media containing 20 uM O-Propargyl Puromycin (MedChemExpress). Cells were stained with the LIVE/DEAD™ Fixable Blue stain (ThermoFisher) according to the manufacturer’s protocol followed by fixation using the Fixation/Permeabilization kit (BD Biosciences). After fixation, cells were washed and permeabilized using 1X perm/wash buffer (BD Biosciences). Cells were stained for OPP using the Click-iT Plus Alexa Fluor 647 Picolyl azide kit (Invitrogen) according to manufacturer’s protocol and using 40% Copper (II) sulfate, and analyzed using a BD-FACS ARIA II.

### Circular RNA Translation Assay

Equal numbers of C57Bl6/J LKS and GMPs were sorted from lineage-depleted bone marrow and cultured as described above. 1 ug of EGFP-circRNA (Creative Biogene, PMCR-0001) was transfected using Lipofectamine MessangerMAXTM (Invitrogen) at a ratio of 1:2 according to manufacturer’s instructions. Media was changed 12 hours post-transfection, and cells were analyzed 24 hours post-transfection. Before analysis, cells were washed with 2% FBS in PBS and resuspended in 2% FBS in PBS containing DAPI for viability. Transfection efficiency was ∼10%.

### Cap-Dependent Translation Inhibition by Small Molecules

Stock solutions of 250 nM Torin-2 (Selleckchem, #S2817) and 25 nM Rocaglamide (MedChemExpress, #HY-19356\CS 5246) were prepared with DMSO. K562 cells stably transduced with the EMCV and RUNX1-bicistronic reporters were treated with and without either drug treatment, and the MFI of GFP and mRFP was calculated one day later on a BD-FACS ARIA II.

### Hair Follicle Immunostaining

Translator mice were sacrificed in second anagen (P29-P30) by CO2 asphyxia. Mouse dorsal skin was dissected and fresh frozen in OCT (Sakura Finetek). 50-µm sections were used for immunofluorescent staining. Slides were fixed for 10 minutes with 4% PFA in PBS at room temperature followed by extensive washes in PBS and 0.3% Tween-20 in PBS. Slides were blocked (5% Donkey serum, 1% BSA, 2% Cold water fish gelatin, and 0.3% Triton in PBS) for 1 – 2 hours at room temperature, incubated with primary antibody overnight at 4°C, washed in 0.3% Tween-20 in PBS, then incubated with secondary antibody for 4 hours at room temperature. Antibodies used: PCAD (goat, R&D AF761, 1:400) and anti-goat IgG 647-conjugated (donkey, Jackson Immuno Research, 705-605-147). Images were acquired using an LSM 880. For evaluating IRES/Cap, the ratio of fluorescent intensity was calculated in HFSC and TACs for GFP and mRFP using FIJI (version 2.30/1.53q). Reporter negative littermates were used to exclude background fluorescence. HFSC and TACs were identified based on PCAD staining and position in hair follicle.

### Production of lentiviral particles encoding bicistronic reporters

The pEF1-EMCV reporter construct was a generous gift from Shira Weinharten-Gabbay (Broad Institute). The RUNX1-IRES element was synthesized and cloned using Bsu36I and NdeI on the original pEF1-EMCV construct by Synbio (Monmouth Junction, NJ, USA). Constructs for evaluating cryptic promoter activity were generated by performing EagI-HF (NEB, #R3505) and BbsI-HF (NEB, #R3539) restriction digestion on the aforementioned constructs, followed by blunting (NEB, #E1201), and T4 DNA ligation (NEB, M0202), per the manufacturer’s instructions. Lentivirus was prepared by transfecting low passage Hek293T with the transfer backbone, Δ8.9, and VSV-G and concentrated by ultracentrifugation of virus containing supernatants 48-72 h post-transfection.

### Production of lentiviral particles encoding inducible sgPTBP1

The destination vector KOBB was generated by modification of pMJ179 (Addgene, Plasmid #85996). After digestion with Hpa1 (NEB, #R0105) and Not1 (NEB, # 0189) the hU6-cloning site-scRNA sequence (Table S3) was cloned into the linearized plasmid using NEBuilder HiFi DNA Assembly Master Mix (NEB, #E2621). Non- and PTBP1-targeting sgRNA sequences were generated by CRISPick (Broad Institute). The sgRNA sequences were inserted by digestion of KOBB destination vector with BamHI (NEB, #R0136) and Xhol (NEB, #0146), and sgRNAs (Table S3) followed by assembly with NEBuilder HiFi DNA Assembly Master Mix (NEB, #E2621). Lentivirus was prepared as described in “*Production of lentiviral particles encoding bicistronic reporters”*. PTBP1 expression was evaluated by comparing cultured and transduced KH2/iCas9^+/-^ *Translator* mice LKS by qPCR. Briefly, RNA was extracted using a RNeasy Plus Micro Kit (Qiagen, #74034) and cDNA was syntheized using an iScript cDNA synthesis kit (Biorad, #1708891). qPCR reactions were performed using iTaq Universal SYBR Green Supermix (Biorad, # 1725122) on a CFX384 Real-Time System (Bio-Rad). The data were analyzed using the 2^-ΔΔCt^ calculations. *Beta-actin* was used as a housekeeping gene. Primers are listed in Table S3.

### Bone Marrow Transplantations

For all transplantations CD45.1 recipient mice were lethally irradiated with 9.5 Gy total body irradiation (split dose, 12 h apart) 16 hours prior to transplantation. All mice were irradiated in a pie cage (Braintree Scientific) with rotation in a JL Shepherd irradiator.

For Mx-cre inducible knockout of PTBP1, LKS sorted from WT (C57Bl6/J) lineage depleted bone marrow or thawed PTBP1^fl/fl^ and Mx-Cre PTBP1^fl/fl^ bone marrow generously gifted by Dr. Bo Porse (University of Copenhagen) were cultured and transduced with lentiviral reporters encoding pEF1-EMCV bicistronic reporter at a multiplicity of infection of 20. For Cas9-inducible knockout of PTBP1, LKS sorted from KH2/iCas9^+/-^ *Translator* mice were transduced with lentivirus encoding KOBB-sgPTBP1 or KOBB-sgNT (see “*Production of lentiviral particles encoding inducible sgPTBP1)*. 16 h post transduction at a multiplicity of infection of 20, cells were washed with 2% FBS in PBS and resuspended PBS. 20,000 transduced LKS were injected retro-orbitally into recipient mice with 200,000 CD45.1 Sca1-depleted marrow, collected using the Anti-Sca1 MicroBead Kit (#130-123-14, Miltenyi Biotec).

Cas9 activation was induced by 3 intraperitoneal injections of 50 ug/g doxycycline (Sigma Millipore, #D3072) every other day in parallel with administration of water containing 2 mg/mL doxycycline and 10 mg/mL sucrose for 10 days. Lineage depleted bone marrow was analyzed 4 months post-transplantation. PTBP1^fl/fl^ and Mx-Cre PTBP1^fl/fl^ HSPCs were identified based on GFP/mRFP+ expression, whereas KH2/iCas9-PTBP1^-/-^ HSPCs were idenfied based on BFP+ (which ranged from ∼10-25%, depending on the immunophenotype). HSPCs were characterized based on immunophenotyping using the panel of antibodies outlined in “*Flow cytometry analysis and sorting”*, except for in iCAS9-PTBP1^-/-^ HSPCs, which were analyzed with CD105 PE-Cy7 and CD150 BV650 used in place of CD150 PE-Cy7.

For primary competitive transplantations, 200,000 CD45.1 whole bone marrow competitor cells were retro-orbitally co injected with 20 CD45.2 SLAM-LKS sorted from the Translator mouse based on the IRES/Cap ratio (low, mid, and high) into lethally irradiated CD45.1 recipient mice. Every four weeks post-transplantation, peripheral blood was obtained by retro-orbital bleeding using heparinized micro-hematocrit capillary tubes (#22-362-566, Fisherbrand) and depleted of red blood cells by ACK lysis (#18-156-721, Quality Biological). Samples were blocked with CD16/32 Fc (#553142, BD Biosciences) for 15 minutes at 4 °C and then stained with CD19-BV785 (Clone 6D5, #115543, BioLegend), CD45.2-APC (Clone 104, #109814, BioLegend), CD3ε-BUV737 (Clone 145-2C11, #612771, BD), CD45.1-BV421 (Clone A20, #110732, BioLegend), Ly-6G-PE-Cy7 (Clone 1A8, #127618, BioLegend), CD8a-BV570 (Clone 53-6.7, #100739, BioLegend), CD4-AF700 (Clone GK1.5, #100430, BioLegend), B220-BV785 (Clone RA3-6B2, #103246, BioLegend) and CD11b-BV711 (Clone M1/70, #101242, BioLegend). Cells were analyzed using a BD-FACS ARIA II (BD). For secondary bone marrow transplantations, one million total bone marrow cells were collected from primary recipients (16 weeks post-transplant) and transplanted into lethally irradiated CD45.1 recipients.

### Cord Blood Transplantations

Human cord blood was purchased from the Pasquarello Tissue Bank (Dana Farber Cancer Institute). CD34+ cells were purified using the EasySep Human Cord Blood CD34 Positive Selection Kit II (#17896, Stem Cell Technologies), as per the manufacturer’s instructions. CD34+ cells were seeded at the concentration of 5E5 cells/mL in serum-free StemSpan medium (#09650, StemCell Technologies) supplemented with penicillin (#15140-122, Gibco, 100 IU/mL), streptomycin (#15140-122, Gibco, 100 IU/mL), glutamine (#25030081, Gibco, 2 mM), SR-1 (#1967-1, Biovision, 1 µM), UM729 (#72332, Stem Cell Technologies, 500 nM), hSCF (#300-07, Peprotech, 100 ng/mL), hFlt3-L (#300-19, 100 ng/mL), hTPO (#300-18, Peprotech, 20 ng/ml), and hIL-6 (#200-06, Peprotech, 20 ng/mL) in a 5% O2, 5% CO2 humidified atmosphere at 37°C. After 24 h of stimulation, cells were transduced with lentiviral particles for bicistronic reporters at a multiplicity of infection of 20 overnight. 300,000 CD34+ cells were washed with PBS+2% FBS, resuspended in PBS, and injected intravenously into 1.5 Gy sub-lethally irradiated NSG mice. Lineage depleted bone marrow was analyzed four months post-transplantation.

### Caloric Restriction and Fasting

For 30 days, translator mice were fed ad libitum either custom control diet (Research Diets, A11051302B) containing 12% protein, 10% fat, and 78% carbohydrates, which was poorly consumed compared to animal facility chow (Prolab IsoPro RMH 3000, LabDiet, 5P75) containing 26% protein, 14% fat, and 60% carbohydrates. Mice were weighed twice per week and monitored for malnourishment. For fasting experiments, animals were restricted from eating for 24, 48, or 72 hours before sacrifice and HSPC analysis by flow cytometry, as described in “*Flow cytometry analysis and sorting”*. Peripheral blood analysis was performed on an Element HT5 (Heska) from retro-orbital bleeding, as described in “*Generation of the Translator Mouse”*.

### Colony Forming Assays

To compare clonogenicity based on IRES/Cap, equal numbers of murine cells (SLAM-LKS: 65, GMP: 165) were sorted based on the ratio and reconstituted in MethoCult M3434 (#03434, Stem Cell Technologies), as per the manufacturer’s instructions. To compare clonogenicity of WT and Translator marrow, 20,000 total bone marrow cells were reconstituted in MethoCult M3434 (#03434, Stem Cell Technologies), as per the manufacturer’s instructions. Colonies were enumerated by visualization ten days post-seeding. For secondary colony assays, 5,000 cells isolated from primary colony assays were plated in MethoCult M3434 (#03434, Stem Cell Technologies), and colonies were enumerated by visualization ten days post-seeding. For human cultures, 1,000 CD34+CD90+EPCR+CD45RA-cultured cord blood cells (transduced four days prior with lentiviral reporters) were sorted based on the ratio of IRES/Cap and reconstituted in Methocult H4434 (#04434, Stem Cell Technologies). Colonies were enumerated by visualization ten days post-seeding. Colony size was measured by ImageJ (NIH) on images taken by an BX41 phase contrast microscope (Olympus) using an Infinity2 microscopy camera (Lumenera).

### Smart-Seq IV RNA Sequencing

To compare the transcriptome of HSPCs based on their relative use of IRES/Cap, 100 SLAM-LKS, GMP, or MEP per well were sorted based on IRES/Cap from the Translator mouse into 2.6 µl of Lysis Buffer (Takara Bio USA, Inc.) followed by snap-freezing at -80°C in preparation for cDNA synthesis using the SMART-Seq v4 assay. The SMART-Seq v4 assay utilizes the SMART technology, switching mechanism at 5’ end of RNA template, to generate full-length cDNA using 11 cycles. The prepared cDNA was assessed for concentration using the Quant-iT Picogreen dsDNA assay kit (Invitrogen, P7589) on the SpectraMax i3 Multi-Mode Detection Platform (Molecular Devices) and normalized to 0.2 ng/µl prior to library prep. Full-length cDNA was fragmented using the Nextera technology in which DNA is simultaneously tagged and fragmented. Tagmented samples were enriched and indexed using 12 cycles of amplification with PCR primers, which included dual 8 bp index sequences to allow for multiplexing (Nextera XT Index Kit). Excess PCR reagents were removed through magnetic bead-based cleanup using PCRClean DX beads (Aline Biosciences) on a the Biomek FXP Single Arm System with Span-8 Pipettor (Beckman Coulter, A31843). Resulting libraries were assessed using a 4200 TapeStation (Agilent Technologies) and quantified by qPCR (Roche Sequencing). Libraries were pooled and sequenced on the Illumina NextSeq500 high output flow cell using paired-end 75 bp reads. Libraries were prepared using the MANTIS Liquid Handler (Formulatrix) and the Biomek FXP Single Arm System with Span-8 Pipettor (Beckman Coulter, A31843). Full-length cDNA was prepared using the SMART-Seq v4 Ultra Low Input RNA Kit for Sequencing (Takara Bio USA, Inc.) and sequencing libraries prepared using the Nextera XT DNA library preparation kit (Illumina).

### Smart-Seq IV Data Analysis

Reads were mapped to reference genome mm10 with ENSEMBL annotation using STAR version 2.5.4a (*57, 58*). Read counts for individual genes were produced using the unstranded count feature in HTSeq 0.9.1 (*58*). Differential expression analysis was performed using the EdgeR package after normalizing read counts and including only those genes with count per million reads (CPM) greater than 1 for one or more samples (*59*). Differentially expressed genes (DEGs) were defined based on the criteria of minimum 2-fold change in expression value and false discovery rate (FDR) less than 0.05 (Table S1). DEGs with logFC greater than 1 or less than -1 with P-Value lower than 0.05 were analyzed for MSigDB Hallmark 2020 using Enrichr (*60–62*).

### Polyvinyl-Alcohol SLAM-LKS Expansion Cultures

SLAM-LKS from lineage depleted Translator mouse bone marrow were sorted based on the ratio of IRES/Cap as single-cells and cultured as described before (*63*). Briefly, PVA expansion media was prepared with F-12 medium (Gibco) supplemented with 10 mM HEPES (Gibco), 1× PSG (penicillin -streptomycin-glutamine Gibco), 1× ITSX (Insulin–transferrin–selenium–ethanolamine Gibco), 1 mg/ml PVA (Polyvinyl alcohol (PVA; 87–90%-hydrolyzed; Sigma), 100 ng/ml AF-TPO (Recombinant animal-free murine thrombopoietin, Peprotech, AF-315-14), and 10 ng/ml AF-SCF (Recombinant animal-free murine stem cell factor, Peprotech, AF-250-03). 20 cells were indexed sorted into a 96 well plate pre-coated with human fibronectin (EMD Millipore) using BD-FACS ARIA II and a 100um nozzle. After three and six days of culture, wells were imaged using a BX41 phase contrast microscope (Olympus) using an Infinity2 microscopy camera (Lumenera). Wells containing megakaryocytes were enumerated visually.

### Generation of CRISPRi cell lines

K562 leukemic cells were first transduced with a lentiviral vector expressing dCas9-BFP-KRAB (Addgene, #135448) and purified for BFP+ cells by flow cytometry. Next, cells were transduced with the bicistronic fluorescent reporter that readouts EMCV-IRES/Cap, and BFP+mRFP+GFP+ cells were sorted as single cells into a 96 well plate for optimal clone selection using a BD-FACS ARIA II. The activity of CRISPRi clone was validated by transducing with positive control sgRNAs targeting ST3GAL3 and SEL1L followed by qPCR analysis of gene expression. sgRNA targeting Gal4 was used as a control.

### Amplification of CRISPRi pooled libraries

Human genome-wide CRISPRi-v2 libraries (Addgene, #83969) were obtained from Addgene. ∼200ng pooled libraries were transformed into MegaX DH10B cells (Thermo Fisher Scientific, C640003) by electroporation (setting: 2.0 kV, 200 ohms, 25 µF in 0.1cm cuvette) to achieve a transformation efficiency of >1,000 colonies/sgRNA. Colonies were grown on LB agar plates overnight and harvested on the next day, followed by plasmid extraction using the NucleoBond Xtra Maxi kit (Macherey-Nagel, 740424). Next, the sgRNA target regions from the amplified libraries were PCR amplified using primers with Illumina adaptors and sequenced by next-generation sequencing to determine the sgRNA distribution of libraries.

### Genome-wide CRISPRi screen

The CRISPRi-v2 libraries were transduced in triplicate into K562 CRISPRi cells in T175 flasks at the MOI ∼0.3 (virus volume was determined by conducting a small-scale pilot test). Two days after transduction, cells were selected by 1 µg/ml puromycin for 5 days. Then, ∼2x107 cells from low, mid, and high IRES/Cap gates were harvested from each triplicate using a BD-FACS ARIA II. Genomic DNA was isolated using NucleoSpin Blood Maxi kit (Macherey-Nagel, 740950). Next, sgRNA target regions were PCR amplified with Illumina adaptors and indexes using Ultra II Q5 (NEB Biolabs, M0544X) and products were purified with DNA Clean & Concentrator-100 (Zymo Research, D4030). sgRNA target regions (260-280bp) were further enriched by gel extraction (Qiagen, 28704) after running agarose gel. Purified PCR products were measured by Qubit and analyzed by TapeStation before sending for next-generation sequencing on Illumina NextSeq 2000.

### CRISPRi screen data analysis

The CRISPR screening data are processed and analyzed using the MAGeCK algorithms (Table S2). Raw sequencing data are pre-processed by using MAGeCK to obtain the read counts for each sgRNA, then MAGeCK robust rank aggregation (RRA) method is used to identify negative and positive selected genes. Briefly, MAGeCK constructed a linear model to compute the variance of guide RNA (gRNA) read counts and assessed the difference in gRNA abundance between low and high IRES/Cap subpopulations. The selection of genes is evaluated from the rankings of gRNAs (by their P-values) using the α-RRA algorithm. For each gene, α RRA assigned P-values for both positive and negative selection. Genes with significant gRNAs were analyzed with Enrichr for KEGG terms (*60–62*).

### Quantification and statistical analysis

GraphPad PRISM 10 was used to plot data and run statistical analysis. Sample sizes were based on prior similar work without the use of additional statistical estimations. All measurements were performed on independent biological replicates unless indicated otherwise.

## Acknowledgments

We extend our sincere gratitude to Dr. Shira Weingarten-Gabbay from the Broad Institute for generously gifting us with the lentiviral EMCV bicistronic construct and for the helpful discussions about IRES biology.

Additionally, we thank Dr. Bo Porse and Dr. Anne-Katrine Frank for supplying us with PTBP1^fl/fl^ BM. This work could not have been accomplished without the contributions of the Harvard Stem Cell Institute Flow Cytometry Core, LASE Biotechnologies, the Harvard Bauer Core, the Harvard Genome Modification Facility, and Animal Facility at Center for Comparative Medicine at MGH.

## Author contributions

DTS and MM conceptualized the study, acquired funding, and wrote the original draft, with DBS, VGS, and PI also contributing to conceptualization. MM, along with LPW and MF, curated the data. Formal analysis was conducted by MM, RIS, and LPW. Methodology was developed by DTS, YCH, VGS, MM, TZ, KG, ESS, MF, SJK, GS, MH, YK, TF, YS, JM, and HK. Investigation was carried out by MM, TZ, AK, TK, KG, ESS, SF, ERA, GS, MH, YK, TF, YS, SK, AS, JM, and HK. MM, MF, ESS were responsible for visualization. Project administration was managed by RIS, MM, and MS. DTS, VGS, DBS, PI, MF, and SJK provided supervision. The manuscript was reviewed and edited by DTS, MM, and DBS

## Data and materials availability

Materials are available, subject to material transfer agreement requests submitted to D. T. S. RNA-sequencing data are deposited at GEO (GSE277603). All other data available in the manuscript or supplementary materials are available from corresponding author upon reasonable request

