## Supplement for "Differentiation stage-specific use of cap-independent and cap-dependent translation initiation in hematopoiesis"

Michael Mazzola *et al.*

**This PDF file includes:**

Figs. S1 to S4
Table S3

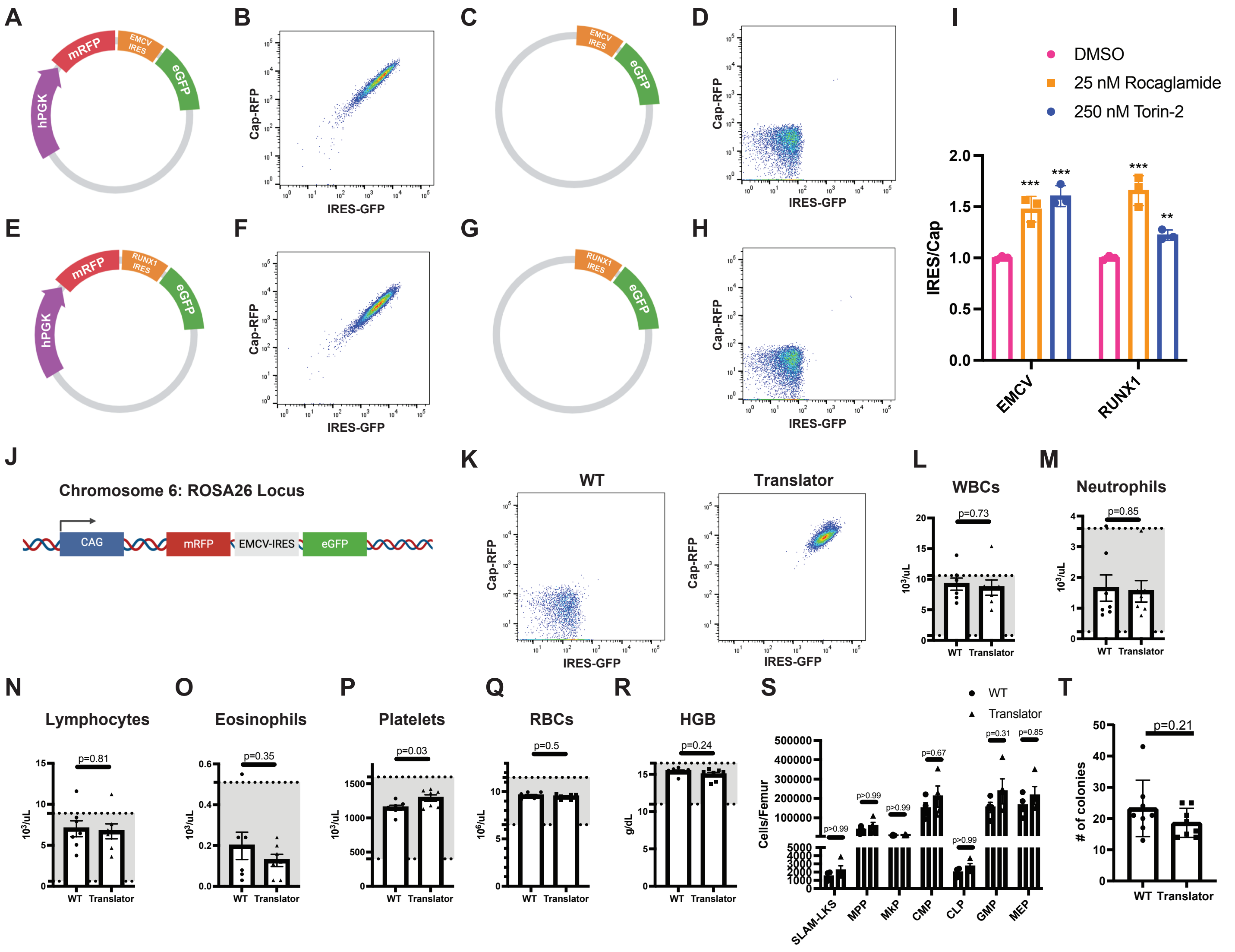


**Fig. S1. Bicistonric fluorescent reporters simultaneously readout rates of cap-dependent and IRES-mediated translation in the Translator mouse, which has comparable hematopoiesis to wildtype littermates.**

**(A)** Graphical representation of bicistronic reporter that simultaneously readout rates of cap-dependent translation (mRFP) and EMCV-IRES cap-independent translation (eGFP). Assisted by BioRender.

**(B)** Flow cytometry flow plot showing co-expression of mRFP and eGFP on K562 cells transduced with lentivirus encoding the bicistronic reporter depicted in (A).

**(C)** Graphical representation of lentiviral vector used to evaluate the EMCV-IRES for cryptic promoter activity. Assisted by BioRender.

**(D)** Flow cytometry flow plot showing no expression of mRFP or eGFP in K562 cells transduced with lentivirus encoding the bicistronic reporter depicted in **(C)**.

**(E)** Graphical representation of bicistronic reporter that simultaneously readout rates of cap-dependent translation (mRFP) and RUNX1-IRES cap-independent translation (eGFP). Assisted by BioRender.

**(F)** Flow cytometry flow plot showing no expression of mRFP or eGFP in K562 cells transduced with lentivirus encoding the bicistronic reporter depicted in **(E)**.

**(G)** Graphical representation of lentiviral vector used to evaluate the RUNX1-IRES for cryptic promoter activity. Assisted by BioRender.

**(H)** Flow cytometry flow plot showing no expression of mRFP or eGFP in K562 cells transduced with lentivirus encoding the bicistronic reporter depicted in **(G)**.
**(I)** K562 cells expressing the EMCV-reporter from **(A)** or the RUNX1-reporter from **(E)** were treated with 25 nM Rocaglamide or 250 nM Torin-2 for 24 h, and the mean-fluorescence intensity was calculated by flow cytometry to evaluate IRES/Cap. Data are normalized to the average of DMSO only controls (n=3).
**(J)** Graphical representation of genomic locus of the *Translator* mouse. Assisted by BioRender.
**(K)** Flow cytometry flow plot depicting co-expression of mRFP and eGFP in *Translator* mouse peripheral blood but not in that of WT littermates.
**(L)** White blood cell (WBC) counts in peripheral blood isolated from *Translator* mice and WT littermate controls. Gray shading represents the normal range (n=7).
**(M)** Neutrophil counts in peripheral blood isolated from *Translator* mice and WT littermate controls. Gray shading represents the normal range (n=7).
**(N)** Lymphocyte counts in peripheral blood isolated from *Translator* mice and WT littermate controls. Gray shading represents the normal range (n=7).
**(O)** Eosinophil counts in peripheral blood isolated from *Translator* mice and WT littermate controls. Gray shading represents the normal range (n=7).
**(P)** Platelet counts in peripheral blood isolated from *Translator* mice and WT littermate controls. Gray shading represents the normal range (n=7).
**(Q)** Red blood cell (RBC) counts in peripheral blood isolated from *Translator* mice and WT littermate controls. Gray shading represents the normal range (n=7).
**(R)** Hemoglobin (HGB) counts in peripheral blood isolated from *Translator* mice and WT littermate controls. Gray shading represents the normal range (n=7).
**(S)** Quantification of HSPCs per femur in 10-week-old WT or *Translator* mice (n=4).
**(T)** Colony formation of BM isolated from 10-week-old WT or *Translator* mice (n=4, two technical replicates).

Data show individual replicates and mean ± SEM. ** $P \leq 0.01$, *** $P \leq 0.01$. Bar graphs represent mean ± SEM. Data were analyzed using a two-way ANOVA with a Sidak's multiple comparisons test **(I)**, a Student's t test **(L-S, U)**, and a one-way ANOVA **(T).**

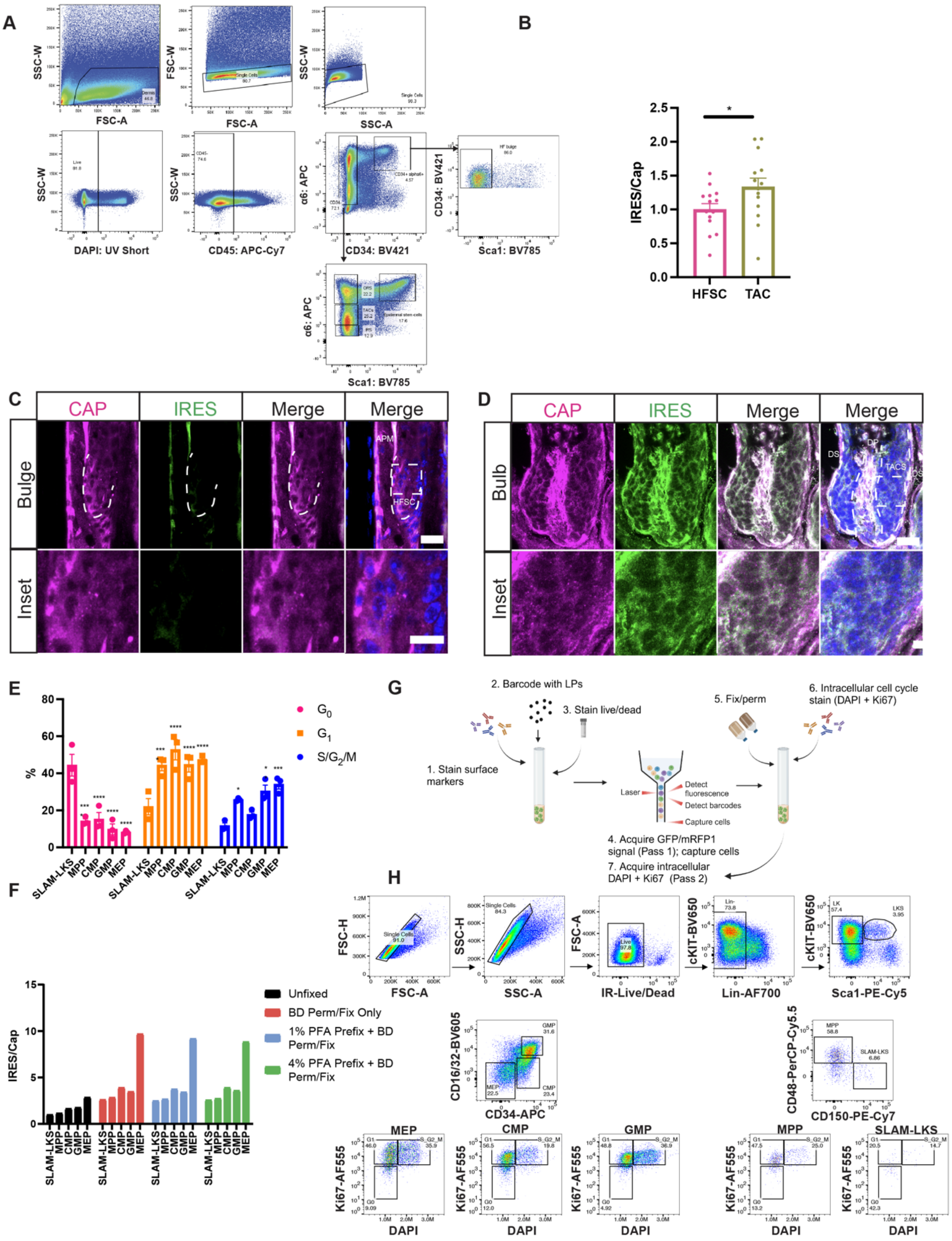
A
B
C
D
E
F
G
H
HFSC
TAC
IRES/Cap
CAP
IRES
Merge
Bulge
Inset
Bulb
G0
G1
S/G2/M
Unfixed
BD Perm/Fix Only
1% PFA Prefix + BD Perm/Fix
4% PFA Prefix + BD Perm/Fix
1. Stain surface markers
2. Barcode with LPs
3. Stain live/dead
4. Acquire GFP/mRFP1 signal (Pass 1); capture cells
5. Fix/perm
6. Intracellular cell cycle stain (DAPI + Ki67)
7. Acquire intracellular DAPI + Ki67 (Pass 2)
Laser
Detect fluorescence
Detect barcodes
Capture cells
SLAM-LKS
MPP
CMP
GMP
MEP
DAPI
Ki67-AF555

**Fig. S2. Hair follicle and cell-cycle analyses.**

**(A)** Representative flow plots to evaluate IRES/Cap in *Translator* mouse hair follicles.
**(B)** IRES/Cap in *Translator* mouse hair follicles, quantified based on immunofluorescence images of the **(C)** HFSCs and **(D)** TACS. DS, dermal sheath. DP, dermal papillae. APM, arrector pili muscle. Error bar (top) 25 µm, error bar (bottom) 10 µm. Blue represents DAPI nuclear staining. (n=14)
**(E)** % HSPCs in $G_0$, $G_1$, and $S/G_2/M$ (n=3). Significance relative to SLAM-LKS.
**(F)** IRES-GFP/Cap-RFP measured in *Translator* mouse HSPCs with various fixation strategies relative to unfixed SLAM-LKS (n=1).
**(G)** Experimental schema for cell cycle analysis of *Translator* mouse HSPCs using a single-cell optical barcoding approach. *Translator* mouse HSPCs were stained for cell-surface markers and labeled with semiconductor laser particles (LPs). Cells were analyzed by flow cytometry to evaluate mRFP, eGFP, HSPC immunophenotypes, and viability. Cells were recollected, fixed and permeabilized, and stained for Ki67 and DAPI. Cells were reanalyzed by flow cytometry. Data from the sequential acquisitions were matched based on the unique LP barcodes to link IRES/Cap and immunophenotype to cell cycle status.
**(H)** Representative flow plots to evaluate IRES/Cap in *Translator* mouse HSPCs using Ki67 and DAPI to define cell cycle status and spectral barcoding to evaluate eGFP and mRFP MFI prior to fixation. Ki67+ cells were gated based on an isotype control.

Data show individual replicates and mean ± SEM. Significance was assessed using an unpaired T-test **(B)** and Dunnett's 2way ANOVA **(E)**. $p \leq 0.05$, $**p \leq 0.01$, $***p \leq 0.001$, $****p \leq 0.0001$.

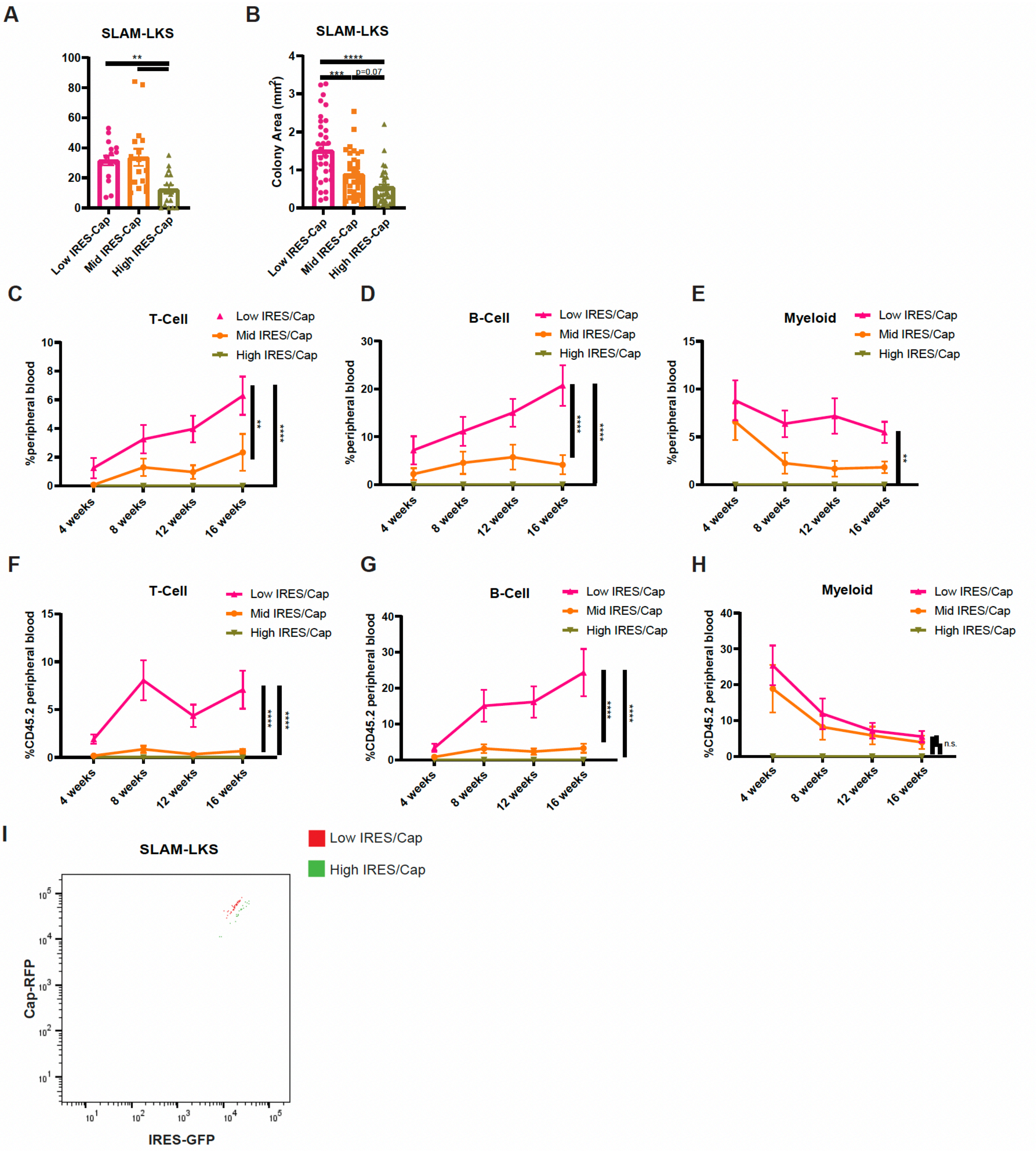
A
SLAM-LKS
**
Low IRES-Cap
Mid IRES-Cap
High IRES-Cap
B
SLAM-LKS
Colony Area (mm2)
****
***
p=0.07
C
T-Cell
%peripheral blood
Low IRES/Cap
Mid IRES/Cap
High IRES/Cap
4 weeks
8 weeks
12 weeks
16 weeks
D
B-Cell
E
Myeloid
F
T-Cell
%CD45.2 peripheral blood
G
B-Cell
H
Myeloid
n.s.
I
SLAM-LKS
Low IRES/Cap
High IRES/Cap
Cap-RFP
IRES-GFP

**Fig. S3. Secondary colony assays and competitive transplantations comparing low, mid, and high IRES/Cap SLAM-LKS clongencity and lineage output.**

**(A-B)** Serial replating of *Translator* SLAM-LKS sorted based on their relative use of IRES/Cap. Colonies were harvested on day 7 and replated in equal numbers. Colonies were then **(A)** enumerated and **(B)** size ($mm^2$) was calculated again on day 14 (n=6).
**(C)** % of live peripheral blood cells that are T-cells derived from donor low, mid and high IRES/Cap *Translator* SLAM-LKS transplanted to primary recipients (n=12).
**(D)** % of live peripheral blood cells that are B-cells derived from donor low, mid and high IRES/Cap *Translator* SLAM-LKS transplanted to primary recipients (n=12).
**(E)** % of live peripheral blood cells that are myeloid-cells derived from donor low, mid and high IRES/Cap *Translator* SLAM-LKS transplanted to primary recipients (n=12).
**(F)** % of live peripheral blood cells that are T-cells derived from donor low, mid and high IRES/Cap *Translator* SLAM-LKS transplanted to secondary recipients (n=12).
**(G)** % of live peripheral blood cells that are B-cells derived from donor low, mid and high IRES/Cap *Translator* SLAM-LKS transplanted to secondary recipients (n=12).
**(H)** % of live peripheral blood cells that are myeloid-cells derived from donor low, mid and high IRES/Cap *Translator* SLAM-LKS transplanted to secondary recipients (n=12).
**(I)** Representative flow cytometry plot that depicts mRFP and GFP for index-sorted low and high IRES/Cap *Translator* SLAM-LKS.

Data show mean ± SEM. ∗$p \leq 0.05$, ∗∗$p \leq 0.01$, ∗∗∗$p \leq 0.001$, ∗∗∗∗$p \leq 0.0001$. Signfance was assessed using a Dunnett's One-way ANOVA **(A-B)** or a two-way ANOVA **(C-H)**.

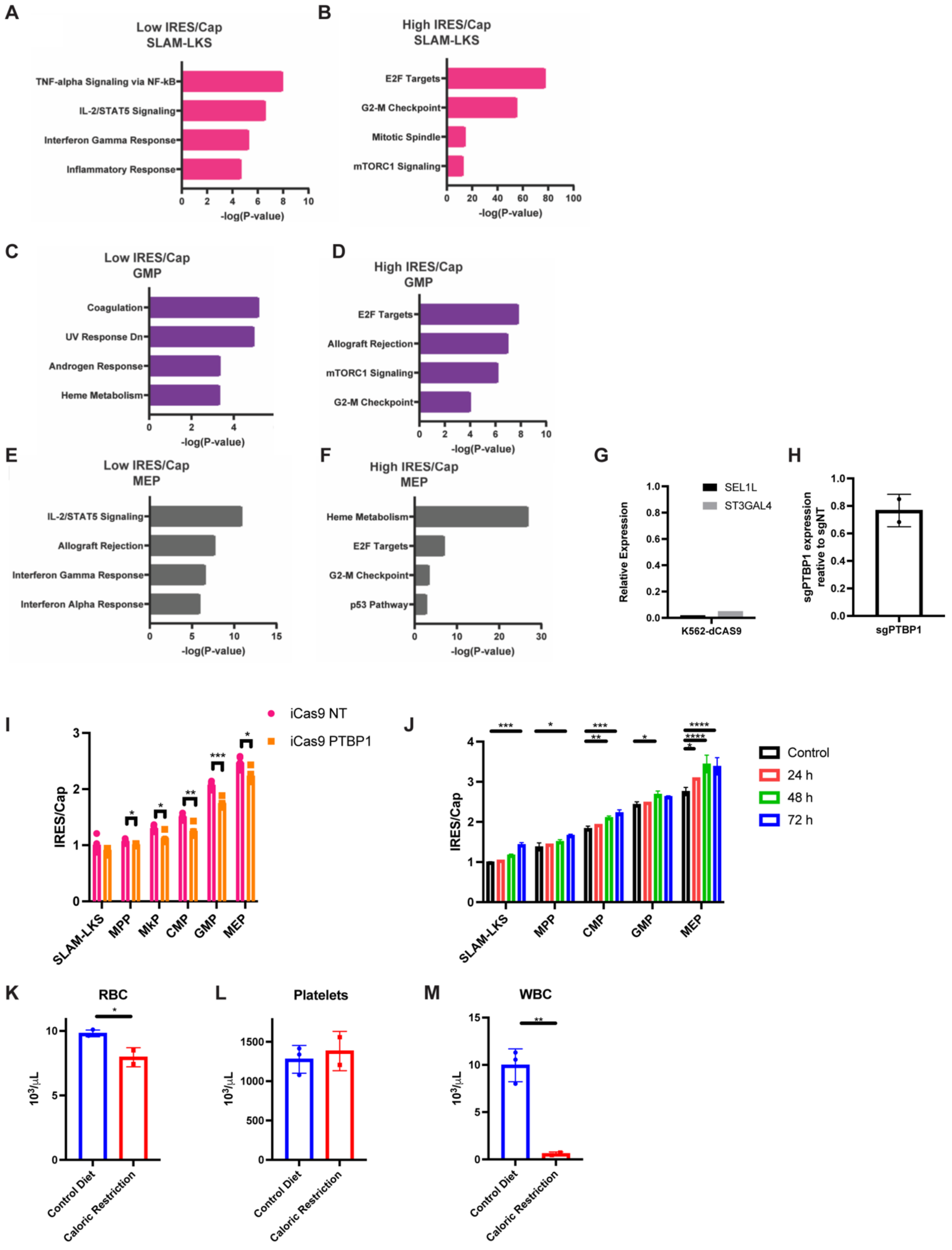
A
Low IRES/Cap
SLAM-LKS
TNF-alpha Signaling via NF-kB
IL-2/STAT5 Signaling
Interferon Gamma Response
Inflammatory Response
-log(P-value)
B
High IRES/Cap
SLAM-LKS
E2F Targets
G2-M Checkpoint
Mitotic Spindle
mTORC1 Signaling
-log(P-value)
C
Low IRES/Cap
GMP
Coagulation
UV Response Dn
Androgen Response
Heme Metabolism
-log(P-value)
D
High IRES/Cap
GMP
E2F Targets
Allograft Rejection
mTORC1 Signaling
G2-M Checkpoint
-log(P-value)
E
Low IRES/Cap
MEP
IL-2/STAT5 Signaling
Allograft Rejection
Interferon Gamma Response
Interferon Alpha Response
-log(P-value)
F
High IRES/Cap
MEP
Heme Metabolism
E2F Targets
G2-M Checkpoint
p53 Pathway
-log(P-value)
G
SEL1L
ST3GAL4
Relative Expression
K562-dCAS9
H
sgPTBP1 expression reative to sgNT
sgPTBP1
I
iCas9 NT
iCas9 PTBP1
IRES/Cap
SLAM-LKS
MPP
MkP
CMP
GMP
MEP
J
Control
24 h
48 h
72 h
IRES/Cap
SLAM-LKS
MPP
CMP
GMP
MEP
K
RBC
10³/μL
Control Diet
Caloric Restriction
L
Platelets
10³/μL
Control Diet
Caloric Restriction
M
WBC
10³/μL
Control Diet
Caloric Restriction

**Fig. S4. Smart-Seq IV hallmark analysis, PTBP1 iCAS9 transplantations, and caloric restriction and fasting data.**
**(A)** Top Hallmark 2020 terms for DEGs enriched in low IRES/Cap SLAM-LKS compared to high IRES/Cap SLAM-LKS (n=3).
**(B)** Top Hallmark 2020 terms for DEGs enriched in high IRES/Cap SLAM-LKS compared to low IRES/Cap SLAM-LKS (n=3).
**(C)** Top Hallmark 2020 terms for DEGs enriched in low IRES/Cap GMP compared to high IRES/Cap SLAM-LKS (n=3).
**(D)** Top Hallmark 2020 terms for DEGs enriched in high IRES/Cap GMP compared to low IRES/Cap SLAM-LKS (n=3).
**(E)** Top Hallmark 2020 terms for DEGs enriched in low IRES/Cap MEP compared to high IRES/Cap SLAM-LKS (n=3).
**(F)** Top Hallmark 2020 terms for DEGs enriched in high IRES/Cap MEP compared to low IRES/Cap SLAM-LKS (n=3).
**(G)** qPCR analysis of K562 CRISPRi cells transduced with SEL1L or ST3GAL4 gRNA relative to transduced with GAL4 gRNA (n=2).
**(H)** Expression of PTBP1 normalized to B-actin in KH2/iCas9 *Translator* LKS transduced with sg-PTBP1 or sg-NT prior to transplantation (N=2).
**(I)** IRES/Cap analysis of HSPCs derived from KH2/iCas9 *Translator* LKS transduced with sg-PTBP1 (n=5) or sg-NT lentivirus (n=6).
**(J)** Analysis of IRES/Cap in HSPCs of *Translator* mice that fasted for 24 (n=1), 48 (n=3), or 72 h (n=2) normalized to control, (i.e. no fasting).
**(K)** Red blood cell (RBC) counts from peripheral blood from *Translator* mice fed control diet (n=3) or those that underwent caloric restriction (n=2).
**(L)** Platelet counts from peripheral blood from *Translator* mice fed control diet (n=3) or those that underwent caloric restriction (n=2).
**(M)** WBC counts from peripheral blood from *Translator* mice fed control diet (n=3) or those that underwent caloric restriction (n=2).

Data show mean ± SEM. $*p \leq 0.05$, $**p \leq 0.01$, $***p \leq 0.001$, $****p \leq 0.0001$. Significance was assessed using multiple T-tests **(I, J, K-M)** and a Dunnett's One-way ANOVA **(K-M)**.

**Table S1. Differential gene expression analysis of SLAM-LKS, GMP, and MEP populations with varying IRES/Cap ratios using Smart-seq V4**

**Table S2. CRISPRi mageck analysis to identify positive and negative regulators of IRES/Cap**

**Table S3. Primers and Cloning**

| **Purpose** | **Sequence** | **Source** |
|---|---|---|
| hU6-cloning site-scRNA | attaggtccctcgacgttaacGAGGGCCTATTTCCCATGATTCCTTCATATTTGCATATACGATACAAGGCTGTTAGAGAGATAATTGGAATTAATTTGACTGTAAACACAAAGATATTAGTACAAAATACGTGACGTAGAAAGTAATAATTTCTTGGGTAGTTTGCAGTTTTAAAATTATGTTTTAAAATGGACTATCATATGCTTACCGTAACTTGAAAGTATTTCGATTTCTTGGCTTTATATATCTTGTGGAAAGGACGAAACACCGgctcgagtactaggatccatgtttAagagctaTGCTGgaaaCAGCAtagcaagttTaaataaggctagtccgttatcaacttgaaaaagtggcaccgagtcggtgctttttttgtggataaccgtattaccgc | Azenta Life Sciences and internal cloning |
| sgPTBP1 | GTGGAAAGGACGAAACACCGGGTTAACTACTATACATCGGGTTTAAGAGCTATGCTGGAA | Azenta Life Sciences |
| sgNT | GCGAGGTATTCGGCTCCGCG | Azenta Life Sciences |
| PTBP1_qPCR_fw | CACCGCTTCAAGAAACCAGGCT | MGH DNA Core |
| PTBP1_qPCR_rev | GTTGCTGGAGAAGAGGCTCTTG | MGH DNA Core |
| Bactin_qPCR_fw | CTCTGGCTCCTAGCACCATGAAGA | MGH DNA Core |
| Bactin_qPCR_rev | GTAAAACGCAGCTCAGTAACAGTCCG | MGH DNA Core |
